# Ventral motor thalamic input to prelimbic cortex mediates cost-benefit decision-making in rats

**DOI:** 10.1101/2022.02.11.480170

**Authors:** Bianca Sieveritz, Shannon Hayashi Duke, Jeffery R. Wickens, Gordon W. Arbuthnott

## Abstract

Corticostriatal neurons in prelimbic cortex contribute to decisions that require a trade-off between cost and benefit. The ventral motor thalamus sends dense projections to many cortical areas, including the prelimbic cortex. We investigated whether this input from the ventral motor thalamus to prelimbic cortex contributes to cost-benefit decision-making. Optogenetic inhibition of ventral motor thalamic axon terminals in prelimbic cortex biased rats towards a high cost-high benefit option and, in anesthetized rats, decreased neuronal activity in deep layers of prelimbic cortex. Stimulation of ventral motor thalamic nuclei induced a neuronal response in deep layers of prelimbic cortex and simultaneous optogenetic inhibition of layer 1 inhibitory interneurons similarly decreased neuronal activity. Our results indicate that ventral motor thalamic input to prelimbic cortex mediates cost-benefit decision-making.

**Significance Statement:** Our results indicate that ventral motor thalamic input to prelimbic cortex plays a critical role in decisions that require a trade-off between two conflicting reward values. Traditionally, ventral motor thalamic nuclei were primarily associated with motor control, but more recently these thalamic nuclei have been implicated in tasks that require animals to choose between two alternatives. Our results highlight the need to reevaluate the role of the ventral motor thalamic nuclei in cognition. Furthermore, prelimbic cortex and, more generally, prefrontal cortex have been associated with chronic stress and major depressive disorder, highlighting the possibility that ventral motor thalamic nuclei might be involved in these disorders.

## Introduction

Prefrontal cortical areas are involved in a variety of decisions that require a trade-off: anterior cingulate cortex regulates the willingness to expend physical (Walton et al., 2003) or mental effort to receive a larger reward (Hosking et al., 2014); orbitofrontal cortex is necessary in risk- and delay-based decision-making (Mobini et al., 2002); dorsomedial prefrontal corticostriatal neurons encode approach-avoidance behavior (Loewke et al., 2021); and prelimbic corticostriatal neurons mediate the trade-off between a more costly, more beneficial and a less costly, less beneficial option (Friedman et al., 2015). Recent studies have highlighted the importance of rodent ventral motor thalamic nuclei (ventromedial, ventral anterior and ventrolateral thalamic nucleus; MT) in tasks that require animals to choose between two options (Guo et al., 2017; Gaidica et al., 2018; Catanese and Jaeger, 2021). In rodents, MT sends dense projections to prelimbic cortex (Herkenham, 1979; Arbuthnott et al., 1990). Taken together, these pieces of evidence raise the question whether MT input to prelimbic cortex is involved in decisions that require a trade-off.

A previous study showed that optogenetic inhibition of prelimbic corticostriatal neurons biased rats towards a more costly, more beneficial option (Friedman et al., 2015). In contrast, the same perturbation did not affect choices made between a high and low benefit, or a high and low cost option (Friedman et al., 2015). We and others previously showed that MT projections to prelimbic cortex preferentially contact corticostriatal pyramidal neurons (Arbuthnott et al., 1990; Collins et al., 2018; Sieveritz and Arbuthnott, 2020). Hence, our first experiment investigated whether MT input to prelimbic cortex is necessary in cost-benefit decision-making. When we inhibited ventral motor thalamic axon terminals in prelimbic cortex on a cost-benefit decision-making task, we found that optogenetic inhibition of MT axon terminals in prelimbic cortex biased rats towards a high cost-high benefit option.

Subsequently, we investigated a potential mechanism for how MT input to prelimbic cortex might induce the observed bias in cost-benefit decision-making. In addition to providing input to prelimbic pyramidal neurons, MT innervates prelimbic layer 1 inhibitory interneurons (Collins et al., 2018; Sieveritz and Arbuthnott, 2020). Layer 1 inhibitory interneurons, in turn, inhibit a network of cortical inhibitory interneurons (Cruikshank et al., 2012), which regulates the activity of deep-layer pyramidal neurons (Jiang et al., 2013; Lee et al., 2015). We found that MT stimulation induced a response in deep-layer pyramidal neurons. Simultaneous inhibition of either MT axon terminals or layer 1 inhibitory interneurons in prelimbic cortex reduced the observed response. Our results show that MT input to both prelimbic pyramidal neurons and prelimbic layer 1 inhibitory interneurons modulates the activity of deep-layer pyramidal neurons in prelimbic cortex. We suggest that inhibition of MT input reduces the activity of deep-layer pyramidal neurons and induces the observed behavioral bias.

## Materials and Methods

### 1. Animals

All applicable international, national, and/or institutional guidelines for the care and use of animals were followed. All procedures performed in studies involving animals were in accordance with the ethical standards of the Okinawa Institute of Science and Technology Graduate University and were approved by the Animal Care and Use Committee of the Okinawa Institute of Science and Technology Graduate University (protocols #2016-131 and #2018-212).

In total, 27 male Sprague-Dawley rats (Charles River Laboratories, Japan) between 4 and 11 weeks of age were used. Rats were single housed in a climate-controlled vivarium, maintained on a 12h light/dark cycle. Rats used in behavioral experiments were housed on a reversed light cycle (lights on at 2200h, lights off at 1000h). Rats used in electrophysiological experiments were either housed on a regular light cycle (lights on at 0700h, lights off at 1900h) or on a reversed light cycle (lights on at 2200h, lights off at 1000h). Food and water were available *ad libitum*. Behavioral experiments were conducted within the last 4 hours of the light cycle or during the dark cycle. Rats were excluded from the study when virus injections extended beyond the target brain region or when the tip of the optical fiber was located outside the target brain area. In addition, rats used in behavioral experiments were excluded from the study if they failed to reach the behavioral criteria specified below.

### 2. The Three Decision-Making Tasks

Twenty-two 4- to 5-week-old Sprague-Dawley rats were trained on a benefit-benefit, cost-cost and cost-benefit decision-making task. Behavioral tasks were modeled after similar tasks presented in another study (Friedman et al., 2015). Rats were trained 6 to 7 days a week and up to 24 days.

In each of the three decision-making tasks rats were presented with a choice between two levers. Each lever was associated with a specific combination of benefit and cost. On the benefit-benefit decision-making task, one lever was associated with a high benefit paired with a low cost, and the other lever was associated with a low benefit paired with a low cost. On the cost-cost decision-making task, one lever was associated with a high cost paired with a high benefit, and the other lever was associated with a low cost paired with a high benefit. On the cost-benefit decision-making task, one lever was associated with a high cost paired with a high benefit, and the other lever was associated with a low cost paired with a low benefit. The high benefit was always 0.1 mL of sweetened condensed milk diluted at 20%. During behavioral training, the low benefit on the benefit-benefit decision-making task was 0.1 mL of sweetened condensed milk diluted at 5%. The first three days of behavioral training on the cost-benefit decision-making task were used to titrate the dilution of sweetened condensed milk for each rat, so that rats would choose both the high cost-high benefit and low cost-low benefit option in roughly 50% of the trials. For the remaining days of behavioral training on the cost-benefit decision-making task and during behavioral testing, the low benefit was 0.1 mL of sweetened condensed milk at the titrated dilution (0-19%). The high cost was a bright light, which was presented for 10 secs at 1.75 kLx. The low cost was a dim light, which was presented for 10 secs at 1.0 1x.

#### 2.1 Apparatus

Behavioral training and testing were conducted in three standard modular test chambers for rats, two of which had a drug infusion top (Med Associates, ENV-008 and ENV-008CT). Each test chamber was enclosed in a custom-made sound-attenuating cubicle and connected to a personal computer by a SmartCtrl connection panel (Med Associates, SG-716B) and a SmartCtrl interface module (Med Associates, DIG-716B). In each test chamber a modified pellet/liquid drop receptacle (Med Associates, ENV-200R3M) was located at the center of the front panel. Two small food trays were attached to the front of the modified receptacle and two blunt needles were placed on top of these small food trays to deliver two different liquids into them. Liquids were delivered using two single speed syringe pumps (Med Associates, PHM-100-3.33) and 50 ml lock type plastic syringes (Terumo). Two retractable levers (Med Associates, ENV-112CM) were placed on each side of the modified receptacle. A white stimulus light (Med Associates, ENV-221M) was mounted above each lever. Stimulus lights were connected to a two level stimulus light fader controller (Med Associates, ENV-226) and could reach a maximum intensity of 200 lx or be reduced to an intensity of 1 lx. Each test chamber was further fitted with a 2900 Hz sonalert module (Med Associate, ENV-223AM), which was mounted in the upper left corner of the front response panel. A 15W LED light (12VMonster, P-15W-E27-CW-12V85V) was placed above each test chamber in a 12V lamp holder (12VMonster, WIRE-E27-2.1MM-2.5M-BLACK) and connected to a SmartCtrl connection panel (Med Associates, SG-716B) by a custom-made electrical circuit board. The SmartCtrl connection panel was connected to a personal computer by a SmartCtrl interface module (Med Associates, DIG-716B). Chronically implanted LED fiber optics, which emitted 590 nm light (TeleLC-Y-8-250, Bio Research Center Co., Ltd.), were controlled by a wireless receiver (TeleR-1-C, Bio Research Center Co., Ltd.). Prior to each behavioral training session, a wireless receiver was secured to the head of each rat and connected to the implanted LED fiber optic. Each wireless receiver was remote-controlled by an infrared signal using a Teleopto Remote (Bio Research Center Co., Ltd.) and a Teleopto Emitter (Bio Research Center Co., Ltd.). Both were placed behind each test chamber. Teleopto Remotes were connected to a personal computer by a passive TTL connection panel (Med Associates, SG-726-TTL) and a SuperPort TTL output interface module (Med Associates, DIG-726TTL-G). The hardware was operated using the Med-PC IV software suite (Med Associates, SOF-735).

#### 2.2 Habituation and Lever Pressing Training

Rats were put on a reversed light cycle (lights on at 2200h, lights off at 1000h) at least 7 days prior to the start of behavioral training, and were handled by the experimenter for at least 3 days out of the 5 days prior to the start of behavioral training. Rats were habituated to 20% sweetened condensed milk diluted in tap water prior to the start of behavioral training. For each rat, 20 mL of 20% sweetened condensed milk diluted in tap water was placed in the home cage on 2 consecutive days out of the 5 days prior to the start of behavioral training.

First, we trained rats to press one of two retractable levers using a modified continuous reinforcement schedule. Each training session started with a 5-second-long tone stimulus (2900 Hz, 65db) and 5 secs later one of the two retractable levers was presented to the rat. Each lever press resulted in retraction of the lever, delivery of 0.1 mL of 20% sweetened condensed milk diluted in tap water and presentation of the stimulus light above the lever at reduced intensity (1 lx) for 10 secs. Once the stimulus light was turned off, a 10-second-long pause occurred. Next, we again presented a 5-second-long tone stimulus (2900 Hz, 65db) to the rat and after another 5 secs the same retractable lever was again presented to the rat. Rats were presented with one 40-minute-long training session each day. In each training session rats were presented with only one lever. The presented lever was alternated each day, unless rats had previously reached criterion on one of the two levers, i.e. performed at least 30 lever presses within 40 mins. If rats had previously reached criterion on one of the two levers, on subsequent days rats were solely trained on the lever that they had not yet reached criterion on. For each of the two levers rats had a total of 6 days to reach criterion, i.e. a maximum of 12 days in total. If rats did not press the presented lever at least one time on two consecutive days, the behavior of rats was shaped by the experimenter. The rat’s paw was placed on the lever and moved downwards to press the lever. Afterwards the rat was moved to the food tray to collect the reward. Shaping was performed for 5 consecutive lever presses. Rats reached criterion on the lever pressing training in an average of 5.1 +/- 2.3 days (mean +/- standard deviation).

#### 2.3 Behavioral Training on the Three Decision-Making Tasks

Once rats had reached criterion on both levers in the lever pressing training, they were trained on the benefit-benefit, cost-cost and cost-benefit decision-making tasks. Rats were always first trained on the benefit-benefit, then on the cost-cost, and last on the cost-benefit decision-making task. Rats completed 40 trials/day on the decision-making task that they were currently trained on. In the benefit-benefit decision-making task, each trial started with a 5-second-long tone stimulus (2900 Hz, 65 dB). In the cost-cost and cost-benefit decision-making task, each trial started with a 5-second-long tone stimulus paired with a 100-millisecond-long bright light flash (1.75 kLx). 5 secs after presentation of the tone stimulus ended, two retractable levers were presented to the rat for up to 10 secs or until a response was made. If no response was recorded within 10 secs, both levers were retracted, and the trial was recorded as omitted. Inter-trial intervals between trials were 25 secs.

Rats were trained on each of the three decision-making tasks for at least 3 days and until criterion was reached. Rats that did not reach criterion after 6 days of training on the benefit-benefit and cost-benefit decision-making task, or after 8 days of training on the cost-cost decision-making task were excluded from the experiment. Criterion on all three decision-making tasks was that less than 20% of trials were omitted. In addition, in the benefit-benefit decision-making task rats had to choose the high benefit option and in the cost-cost decision-making task the low cost option on at least 52% of the trials. Overall, 53.7% of rats learned the task. Reported animal numbers only include rats that reached criterion and were included in the data analysis. Across rats that learned all three decision-making tasks, criterion was reached in an average of 3.3 +/- 0.7 days on the benefit-benefit decision-making, in an average of 4.3 +/- 1.7 days on the cost-cost decision-making task, and in an average of 2.4 +/- 1.2 days on the cost-benefit decision-making task (mean +/- standard deviation).

We used the first three days of behavioral training on the cost-benefit decision-making task to titrate the dilution of sweetened condensed milk delivered as low reward, so that rats would choose both the high cost-high benefit and the low cost-low benefit option in about 50% of the trials. The dilution of the sweetened condensed milk offered as a low benefit was systematically varied across the 120 trials presented over the first 3 days of behavioral training on the cost-benefit decision-making task. The dilution was reduced every 20 trials starting at 13% followed by 11%, 8%, 5%, 2% and then pure tap water. For each dilution the percentage of trials that the rat chose the low cost-low benefit option out of the overall number trials that were not omitted was calculated and a psychometric curve was fitted to the data to estimate the best dilution of sweetened condensed milk for each rat. The psychometric curve was fitted using the FitWeibull function from the PsychoPy package (Peirce, 2007) under Python 2.7. For each rat, guesses for the threshold and slope of the psychometric curve were made by the experimenter based on the collected data. Usually, the guess for the threshold was equal to the dilution at which rats chose the low cost-low benefit option in about 50% of the trials. The initial guess for the slope was 2, but if no optimal parameters for the psychometric curve were found by the function, the parameter for the slope was increased to 5 and then doubled until optimal parameters were determined. The resulting dilution of sweetened condensed milk was rounded to the nearest whole number. Given that the individual dilution of sweetened condensed milk for each rat was determined during training on the cost-benefit decision-making task, the individual dilution could not be used for initial training on the benefit-benefit decision-making task. Thus, on the benefit-benefit decision-making task all rats were trained with a low benefit of 0.1 ml sweetened condensed milk diluted at 5%.

### 3. Surgeries

Virus injections, LED fiber optics or both were placed in 9- to 11-week-old male Sprague-Dawley rats. Rats were anesthetized with 5% isoflurane delivered with room air (0.5-1 L/min; Small Animal Anesthetizer, Muromachi, MK-A100, Japan). Rats were positioned in a stereotaxic frame (Narishige, SR-5R-HT, Japan) and anesthesia was maintained at 2% isoflurane delivered with room air (0.5-1 L/min). Virus injections were performed with a Hamilton syringe (Neuros Syringe, 0.5 μL, Neuros Model 7000.5 KH, point style 3, Hamilton, 65457-01, United Kingdom or Neuros Syringe, 1.0 μL, Neuros Model 7001 KH, point style 3, Hamilton, 65458-01, United Kingdom) at an injection speed of 100 nL/min and the syringe remained at the target location for 10 min after the injection before it was slowly retracted from the brain. All experiments involving recombinant DNA were approved by the Biosafety Committee of the Okinawa Institute of Science and Technology Graduate University (protocol #RDE-2017-003) and all applicable international, national, and/or institutional guidelines were followed.

For electrophysiological experiments in anesthetized rats, unilateral injections of 50-70 nL AAV5-CAG-ArchT-GFP (titer ≥ 7× 10^12^ vg/mL; gift from Edward Boyden and purchased through UNC Vector Core; now commercially available from Addgene viral preparation #29777-AAV5; http://n2t.net/addgene:29777; RRID:Addgene_29777) were placed in MT (from interaural zero AP +7.76 to +9.97, ML −1.2, from dura DV −6.6; Paxinos and Watson, 2004) or in prelimbic cortical layer 1 (from bregma AP +1.32 mm, ML +0.8 mm with the stereotaxic rotated at a 30° angle towards posterior, from dura DV −3.58 mm with the stereotaxic tilted at a 30° angle; Paxinos and Watson, 2004). Experiments were performed 12-16 days after the virus was injected and rats were perfused after the experiment.

For behavioral experiments, unilateral injections of 50-70 nL of either an adeno-associated virus expressing archaerhodopsin (AAV5-CAG-ArchT-GFP; titer ≥ 7× 10^12^ vg/mL; gift from Edward Boyden and purchased through UNC Vector Core; now commercially available from Addgene viral preparation #29777-AAV5; http://n2t.net/addgene:29777; RRID:Addgene_29777) or a control virus (AAV5-CAG-GFP; titer ≥ 7× 10^12^ vg/mL; gift from Edward Boyden and purchased through UNC Vector Core; now commercially available from Addgene viral preparation #37825-AAV5; http://n2t.net/addgene:37825; RRID:Addgene_37825) were placed in MT (from interaural zero AP +6.31 to +10.26, ML −1.2, from dura DV −6.6; Paxinos and Watson, 2004). In the same aseptic surgery, a short optical fiber attached to a 590 nm light emitting diode (LED fiber optic) was chronically implanted through the contralateral hemisphere into ipsilateral prelimbic cortical layer 1 (from bregma AP +0.898 mm, ML +1.084 mm with stereotaxic turned at a 30° angle towards posterior, from dura DV −3.58 mm with the stereotaxic tilted at a 30° angle; Paxinos and Watson, 2004). Each LED fiber optic had a diameter of 250 μm and emitted 590 nm light (TeleLC-Y-8-250, Bio Research Center Co., Ltd.). LED fiber optics were fixed to the skull with dental cement (Super-Bond C&B, Sun Medical, Japan). Behavioral testing was conducted after a recovery period of 12-16 days and rats were perfused after 26-28 days.

### 4. Experimental Design and Statistical Analyses

#### 4.1 Experimental Design - Choice Behavior on the Three Decision Making Paradigms

22 male Sprague-Dawley rats were used (Charles River Laboratories, Japan); 10 rats that had been injected with the archaerhodopsin-expressing virus (ArchT rats) and 12 rats that had been injected with the control virus (controls). Rats were between 4 to 5 weeks of age, when we started habituation to the sweetened condensed milk and behavioral training. Rats were 9 weeks of age at the time of surgery and 11 weeks of age, when we started behavioral testing. The required number of animals for behavioral experiments was predicted a priori based on a similar study conducted in the past (Friedman et al., 2015).

Assignment of rats to groups for behavioral training was pseudo-randomized. For the initial lever pressing training we presented the left lever on the first day and the right lever on the second day for one group of rats, while for a second group of rats the right lever was presented on the first day and the second lever was presented on the second day. For behavioral training and testing for one group of rats the left and for a second group of rats the right lever was associated with the high benefit, high cost and high cost-high benefit option. The lever on the opposite side was associated with the other choice option. Assignment of rats to the ArchT or control groups was also pseudo-randomized.

Rats had a 12- to 16-day-long recovery period after surgery before their choice behavior on each of the three decision-making tasks was assessed over a period of 9 days. Rats were presented with 40 trials per day over 3 consecutive days on each decision-making task. The first 20 trials were presented without delivery of the 590 nm light (light OFF) and the last 20 trials were paired with delivery of the 590 nm light (light ON). In ArchT rats, light delivery induced optogenetic inhibition in MT axon terminals located in prelimbic cortical layer 1. In total, we presented rats with 60 light OFF and 60 light ON trials on each decision-making task. The 590 nm light was turned on at the same time as the tone stimulus and remained on until rats made a choice or until the levers were retracted, i.e. a maximum of 20 secs.

The implanted LED fiber optics were controlled by wireless receivers. The maximum light intensity that was reached using the wireless receivers and LED fiber optics was between 0.7-1.2 mW. Given that LED fiber optics had a diameter 250 μm, the emitted 590 nm light reached approximately 3-6 mW/mm^2^.

We decided on a block design for presentation of light OFF and ON trials, since a previous study suggested that sustained optogenetic inhibition of axon terminals may cause unexpected long-term effects such as an increase in spontaneous neurotransmitter release (Mahn et al., 2016). Hence, light ON trials were always presented after light OFF trials. All rats were first tested on the benefit-benefit, then on the cost-cost and last on the cost-benefit decision-making task. We presented the decision-making tasks in this specific order, to increase rats’ retention of the tasks. A timeline for behavioral testing is provided in Figure 1.

**Figure 1.**
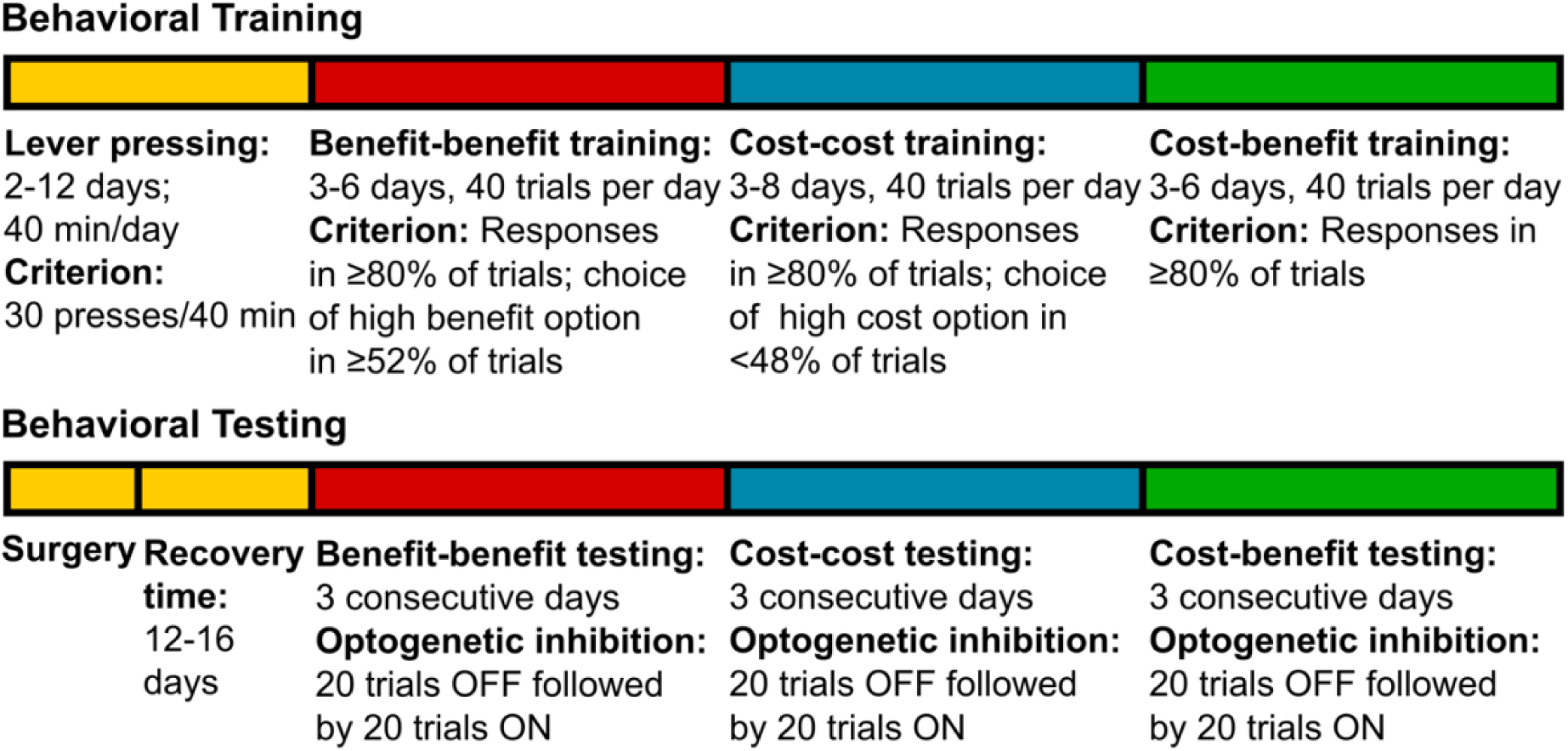
Timeline of behavioral training and testing for behavioral experiments assessing the effect of optogenetic inhibition of MT axon terminals in prelimbic cortical layer 1.

Behavioral responses in each session were stored in an automatically generated text document. We extracted any variables that were analyzed from these text documents using custom-written Python 3.7 scripts. The reaction times stored in each text document represented a measurement from the presentation of the tone stimulus marking the beginning of each trial to the time rats indicated their choice by pressing one of the two levers. However, reaction times were corrected to represent times from presentation of the two levers to the time rats indicated their choice.

#### 4.2 Statistical Analyses - Choice Behavior on the Three Decision Making Paradigms

To quantify and compare decision-making behavior of rats on the three decision-making tasks, we primarily assessed the percentage of trials, in which rats chose 1) the high benefit option on the benefit-benefit decision-making task, 2) the high cost option on the cost-cost-decision-making task or 3) the high cost-high benefit option on the cost-benefit decision-making task out of the total number of non-omitted trials.

In addition, to test whether motor function was disrupted by optogenetic inhibition of MT axon terminals in prelimbic cortical layer 1, we assessed 1) the percentage of omitted trials; and 2) the average reaction time across all non-omitted trials.

To analyze the effect of the injected virus and of light ON versus OFF, we performed separate ANOVAs for each decision-making task and each variable using the injected virus as between and treatment as within animal variable. When we observed a main or interaction effect, we performed two post-hoc paired t-tests to compare the treatment and non-treatment condition within each group of rats. To confirm that effects were not caused by pre-existing differences in behavior between the two groups of rats, we further performed two Student’s t-tests to compare behavior on the first 20 trials of the last day of training as well as behavior in the non-treatment condition between the two groups of rats. To confirm that effects were not caused by surgery, we also compared the behavior pre- and post-surgery within each group of rats. We usually performed 6 t-tests to analyze each behavioral variable. We applied a Bonferroni correction to the significance level resulting in a significance level of p=0.0083. However, when comparing the effect of optogenetic inhibition of MT axon terminals in prelimbic cortical layer 1 on choice behavior, we also compared differences in choice behavior between light ON and OFF trials for each individual day of behavioral testing. In addition, we compared differences in the percentage of high benefit and high cost choices between the first and last 20 trials on the last three days of behavioral training. Overall, the number of statistical tests resulted in an adjusted significance level of p=0.0028.

We observed a significant difference in the percentage of high cost-high benefit choices between light ON and OFF trials for ArchT rats. To quantify the size of the difference, we subtracted the percentage in light OFF from that in light ON trials. We performed a Student’s t-test on this new metric (significance level of p=0.05), comparing it between controls and ArchT rats, to analyze whether the difference in the percentage of high cost-high benefit trials was significantly different between the two groups of rats.

We adjusted individual dilutions of sweetened condensed milk for each animal, so that each animal would choose the high cost-high benefit option as well as the low cost-low benefit option in about 50% of non-omitted trials. We used the first 3 days of behavioral training on the cost-benefit decision-making task to adjust individual dilutions. We systematically varied the dilution of sweetened condensed milk on every 20 trials. Hence, data on the percentage of high cost-high benefit choices, which was collected across these 3 days, does not represent the actual choice behavior of rats on days after we determined and used their individual dilutions. Only a few rats did not reach criterion on the cost-benefit decision-making task within the first 3 days, resulting in few rats being trained for additional days. When running statistical tests that involved data from the last day of behavioral training on the cost-benefit decision-making task, we only used the data from the first 20 trials on the last day of behavioral training for these few rats. For all other rats, when the determined individual dilution corresponded to a dilution that was used during the adjustment process, we used data from those 20 trials. Otherwise, when the individual dilution was between dilutions used during the adjustment process, we averaged data from the 20 trials with the next higher and the 20 trials with the next lower dilution. Given the limitations of this approach, we compared a second metric between groups of rats to determine whether pseudo-random assignment of rats to either the control or ArchT group influenced the percentage of high cost-high benefit choices within each group. We analyzed for each group of rats whether the individual dilution of sweetened condensed milk correlated with the percentage of high cost-high benefit choices in light OFF trials during behavioral testing. We assumed that if these two metrics did not correlate, pseudo-random assignment of rats would not have influenced the percentage of high cost-high benefit choices in each group, even if the determined individual dilutions of sweetened condensed milk would have varied vastly. We calculated Kendall’s Tau to determine the correlation between the two metrics. We chose Kendall’s Tau since individual dilutions of sweetened condensed milk were rounded to the closest whole number and, hence, on an ordinal scale. We chose Kendall’s Tau over Spearman’s rho due to the small sample size and higher robustness of Kendall’s Tau for small sample sizes.

Statistical analyses were performed in R (version 3.6.3). We tested the assumption of the data being normally distributed, which is a prerequisite for parametric tests, with Levene’s test and the assumption of homogeneity of variances with Shapiro’s test.

#### 4.3 Experimental Design – In-vivo Electrophysiology

We acquired extracellular recordings from deep-layer pyramidal neurons in anesthetized, male Sprague-Dawley rats (Charles River Laboratories, Japan). We stimulated MT and, in four rats, simultaneously inhibited MT axon terminals in prelimbic cortical layer 1 using optogenetics. Surgeries were performed when these rats were between 9 to 11 weeks of age, and data was collected when rats were between 11 to 13 weeks of age. We recorded from a total of 39 cells. In three rats, we stimulated MT and simultaneously inhibited prelimbic layer 1 inhibitory interneurons using optogenetics. Surgeries were performed when these rats were between 9 to 10 weeks of age, and data was collected when rats were between 11 to 12 weeks of age. We recorded from a total of 37 cells. The required number of cells for electrophysiological experiments in anesthetized rats was predicted based on previous studies (Yuan et al., 1985; Brecht and Sakmann, 2002; Martin-Cortecero and Nuñez, 2016).

12-16 days after an AAV5-CAG-ArchT-GFP injection had been placed into MT or prelimbic layer 1, rats were anesthetized with 5% isoflurane delivered with room air (1.5-2 L/min, Classic T3 Vaporizer, SurgiVet) and positioned in a stereotaxic frame (custom-build from a Model 1730 Intracellular Frame Assembly from David Kopf Instruments). Anesthesia was maintained at 1.5-2% isoflurane delivered with room air (1.5-2 L/min). An optical fiber with a diameter of 250 μm connected to a 590 nm fiber-coupled LED (ThorLabs, M590F2) that was powered by a high-power, 1-channel LED driver with pulse modulation (ThorLabs, DC2100) was placed in prelimbic layer 1 (from bregma AP +0.898 mm, ML +1.084 mm with stereotaxic turned at a 30° angle towards posterior, from dura DV −3.58 mm with the stereotaxic tilted at a 30° angle; Paxinos and Watson, 2004). A concentric bipolar platinum/iridium microelectrode with a wire diameter of 25 μm (FHC, CBBPC75) was placed in close proximity to MT (from bregma AP −4.40 mm, ML −1.20 mm, from dura −6.98 mm with the stereotaxic tilted at a 20° angle; Paxinos and Watson, 2004) and was connected to a biphasic current stimulator (Digitimer, DS4) to stimulate MT and the ascending projections. Both, the 1-channel LED driver and the biphasic current stimulator, were connected to a personal computer by a low-noise data acquisition system (Molecular Devices, Axon CNS Digidata 1440A) that was controlled by Clampex 10.3 to deliver light pulses for optogenetic inhibition and microstimulation using a previously specified protocol. To perform extracellular recordings from neurons in deep layers of prelimbic cortex, glass electrodes with an impedance between 50-110 MΩ were pulled on a P-1000 micropipette puller (Sutter Instrument, P-1000) and placed in prelimbic cortex in lower layer 2/3, layer 4 or layer 5A. Glass electrodes were filled with an internal solution of 3.0 molar potassium methyl sulfate (KMeSO_4_) that contained goat anti-rat AlexaFluor 594 at a dilution between 1:50 to 1:200, and were coated with 1,1’-dioctadecyl-3,3,3’,3’-tetramethylindocarbocyanine perchlorate stain (DiI stain, 2.5 mg/ml, Thermo Fisher, D282) diluted in one part of ethanol and 9 parts of water. Glass electrodes were connected to a personal computer by a low-noise data acquisition system (Molecular Devices, Axon CNS Digidata 1440A), a Hum Bug Noise Eliminator (Quest Scientific) and a computer-controlled microelectrode amplifier (Molecular Devices, Axon CNS, Axoclamp 900A) and recordings were performed using the Axoclamp 900A software and Clampex 10.3.

Glass electrodes were lowered to the dorsal end of prelimbic cortex in 10 μm steps and then advanced to the ventral end of prelimbic cortex in 0.2-1 μm steps using a micropositioner (David Kopf Instruments, Model 2660). When spiking activity was observed MT and its ascending projections were stimulated (light OFF trials). Multiple iterations of ten 0.5-millisecond-long −5 mA stimulations were delivered at a rate of 10 Hz over a total time period of 1 sec (light OFF trials). Each iteration was followed by a 3-seconds-long pause, before the next iteration of microstimulations began. In rats that had received injections of the archaerhodopsin-expressing virus into MT, ten iterations of microstimulation were delivered. In rats that had received injections of the archaerhodopsin-expressing virus into prelimbic cortical layer 1, five iterations of microstimulations were delivered. Data was digitized at 250000 Hz.

When stimulation of MT and its ascending projections induced spiking activity at the recording site in deep prelimbic layers, MT stimulation was paired with optogenetics. In each case, five iterations of MT stimulations were paired with 20-second-long delivery of 590 nm light to prelimbic cortical layer 1 (light ON trials). In rats that had received virus injections into MT, we ran this protocol twice with a 4-second-long pause between the two runs. In rats that had received virus injections into prelimbic cortical layer 1, we ran this protocol thrice and decreased the strength of stimulations with each run. In the first run stimulations were delivered at −5 mA, in the second run at −3 mA, and in the third run at −1 mA. Data was digitized at 20000 Hz. All data was recorded using a high-band pass filter at 100 Hz and a lowpass Bessel filter at 10 kHz.

#### 4.4 Statistical Analyses - In-vivo Electrophysiology

We used custom-written Matlab scripts (Matlab 2019b) to analyze electrophysiological data collected in anesthetized rats. Data were filtered at 100 Hz to 10 kHz at the time of recordings and no further filtering was done. To analyze the effect of optogenetic inhibition of MT axon terminals or inhibitory interneurons in prelimbic cortical layer 1 on neuronal cluster responses recorded in deep layers of prelimbic cortex, we defined segments starting 20 ms before and ending 80 ms after MT stimulations were delivered. We normalized each data segment by subtracting the average activity measured across the last 40 ms of each segment from each data point in the segment. MT stimulations induced an artefact, which was truncated from any presented figures by removing data from the time of MT stimulation until 2 ms after. For each cell, the threshold to identify extracellular spikes was determined by using the Matlab ‘findpeaks’-algorithm with a minimum spike prominence of 0.5 mV on data segments from light OFF trials. We identified the spike with the largest spike prominence and identified extracellular spikes by running the ‘findpeaks’-algorithm again, but using 30% of the largest identified spike prominence as minimum spike prominence. We constructed a peri-stimulus time histogram with a bin width of 0.1 ms from the data. To compare the average number of spikes per millisecond between light ON and OFF trials within the first 20 ms after MT stimulation, we conducted a Wilcoxon ranked-sign test across cells.

To compare the width of spikes during segments of spontaneous activity, after MT stimulation in light OFF trials and after MT stimulation in light ON trials, we analyzed the last 2 seconds of each light OFF sweep. Each light OFF sweep started with ten MT stimulations followed by a 3-second-long pause before the next sweep started. Spontaneous activity was observed within this 3-second-long pause of which we analyzed the last 2 seconds. Data within these 2-second-long segments were normalized by subtracting the average activity measured across the last 40 ms of each segment from each data point in the segment. The width of spikes within these segments of spontaneous activity, for light OFF and for light ON trials were identified by running the Matlab ‘findpeaks’-algorithm and using the previously determined value for minimum spike prominence. Spike widths were summarized in histograms. In addition, segments showing activity after MT stimulation in light OFF trials as well as spontaneous activity were plotted for selected cells.

To test whether rebound spiking was observed upon turning the 590 nm light off, which deactivated the archaerhodopsin and terminated optogenetic inhibition of MT axon terminals, we extracted segments from light ON trials starting 100 ms before and ending 100 ms after the light was turned off. Data was normalized by subtracting the mean activity across each segment from each data point in the segment. We again identified extracellular spikes by running the Matlab ‘findpeaks’-algorithm and using the previously determined value for minimum spike prominence. We constructed a peri-stimulus time histogram from the data using a bin size of 1 ms. In addition, the average number of spikes per second for the 100 ms before and the 100 ms after the light was turned off was calculated and indicated in the peri-stimulus time histogram by a red dotted line.

### 5. Perfusion

Rats were anesthetized with approximately 10 ml isoflurane delivered on tissue paper and medetomidine hydrochloride (1.0 mg/kg, intraperitoneal), before being transcardially perfused with either 100 mL 0.1 M phosphate buffer with 2 mg/100 mL heparin (4 °C, pH7.4) or 100 mL of 10% sucrose diluted in Milli-Q, both followed by 150 mL Lana’s fixative (room temperature, 4% depolymerized paraformaldehyde and 14% saturated picric acid solution, Sigma Aldrich, P6744-1GA, in 0.16 M phosphate buffer, pH7.4). Brain tissue was post-fixed for at least 36 h and afterward stored in a 50/50 mixture of Lana’s fixative and 20% sucrose in phosphate-buffered saline (PBS) until sectioning, i.e. a minimum of 36 h and up to 2 months.

### 6. Immunohistochemistry

80 μm-thick coronal or sagittal sections that contained prelimbic cortex (interaural zero +11.52 to +13.68; Paxinos and Watson, 2004), striatum (interaural zero +8.40 to +11.28; Paxinos and Watson, 2004) and/or MT (interaural zero +4.80 to +8.76; Paxinos and Watson, 2004) were taken using a freezing microtome (Yamato, REM-710, Japan). For some sections no immunohistochemistry was performed. These sections were washed in PBS for 4 x 5 min before being mounted. When immunohistochemistry was performed, sections were first washed in PBS for 3 x 5 min. Next, sections were incubated in 20% goat serum (Vector Laboratories, S-1000) or, when the primary antibody was grown in goat, in 20% donkey serum (Millipore, S30-100ML) diluted in PBS, containing 0.05% sodium azide and 0.3% Triton X-100 (Sigma Aldrich, 234729-100ML), at room temperature for 1h. Afterward, sections were incubated for 18h-36h at 4°C in primary antibody solution containing primary antibodies diluted in PBS, 0.05% sodium azide and 0.3% Triton X-100. Then, sections were washed in PBS for 3 x 5 min before being incubated at room temperature for 3h in secondary antibody solution containing secondary antibodies diluted in PBS, 0.05% sodium azide and 0.3% Triton X-100. Finally, sections were washed in PBS for 4 x 5 min. Dilutions of primary and secondary antibodies varied depending on the antibodies used and are provided in table 1. All sections were mounted on glass slides in several drops of mounting medium and sealed with a cover slip. We used UltraCruz Hard-set Mounting Medium with DAPI (Santa Cruz, sc-359850) or Vectashield HardSet Mounting Medium with DAPI (Vector Laboratories, H-1500) that also provided DNA counterstaining of cell nuclei. Secondary antibodies were acquired from Millipore or Invitrogen.

**Table 1.** Primary and secondary antibodies used in immunohistochemistry. A summary of dilutions of primary and secondary antibodies in respective antibody solutions, and the dilution of Triton X-100 in primary antibody solutions.

| Primary antibody | Secondary antibody |
| --- | --- |
| Mouse anti-GAD67<br>(Millipore, MAB5406, RRID:AB_2278725):<br>1:1000 dilution, 0.3% Triton X-100,<br>18-36h incubation time | Alexa Fluor 594 goat anti-mouse:<br>1:200 dilution, 3h incubation time |
| Rabbit anti-GFAP<br>(bioSensis, R-1374, RRID:AB_2492897):<br>1:1000 dilution, 0.3% Triton X-100,<br>18-36h incubation time | Alexa Fluor 594 goat anti-rabbit:<br>1:200 dilution, 3h incubation time |

### 7. Microscopy

Histological verification of the placement of virus injections or LED fiber optics was performed with a fixed stage BX51WI upright microscope from Olympus. A 100W mercury lamp was used as a light source. Alexa Fluor 594 was visualized using an excitation filter with a peak at 555 nm and a bandwidth of 25 nm, and an emission filter with a peak at 605 nm and a bandwidth of 25 nm. Images were acquired with a 4x objective with a numerical aperture (NA) of 0.16 with the Neurolucida software (MBF Bioscience).

To acquire the high-resolution images presented in this paper, laser confocal scanning microscopy was performed with a Zeiss LSM 780 microscope at room temperature. Images were taken with a 10x objective with a NA of 0.45 or a 63x oil objective with a NA of 1.46 (both objectives from Zeiss). The 63x objective was used with fluorescence-free Immersol immersion oil 518F from Zeiss. Any tiled overview images were acquired with 10% overlap between tiles and stitched using the stitching algorithm provided by the ZEN 2011 SP7 FP1 software (Zeiss, black edition, version 14.0.8.201). An overview of the excitation and emission peaks of used fluorochromes, the light source and filter set used for excitation, and the emission range used are provided in table 2.

**Table 2.** Used fluorochromes. Overview of the excitation and emission peak of used fluorochromes, the light source and filter set used for excitation, and the emission range used.

| Fluoro-<br>chrome | Excitation | Emission | Light<br>Source | Filter Set | Emission<br>Range |
| --- | --- | --- | --- | --- | --- |
| DAPI | 358 nm | 461 nm | 405-30<br>laser diode | MBS 488/561/633 dichroic beam<br>splitter; MBS -405 dichroic beam<br>splitter | 410-495 nm |
| GFP | 395 nm | 509 nm | Argon<br>laser | MBS 488 dichroic beam splitter | 490-590 nm |
| Alexa<br>Fluor 594 | 590 nm | 617 nm | DPSS 561-10 laser | MBS 458/561 dichroic beam<br>splitter | 585-734 nm |

Images were acquired using the ZEN 2011 SP7 FP1 software (Zeiss, black edition, version 14.0.8.201). Brightness and contrast were adjusted using ZEN 2011 SP7 FP1 software (Zeiss, black edition, version 14.0.8.201) and the “Enhance Contrast” option in Fiji (ImageJ version 1.51n or 1.52p). When necessary, brightness and contrast were adjusted separately for each channel. For final publication, brightness and contrast of images were additionally adjusted using Adobe Photoshop CS5 Extended (version 12.0.4 x64). Adjustments to brightness and contrast were always applied equally across the entire image.

### 8. Code and Data Availability

Raw behavioral data, raw electrophysiology data and analysis scripts used to generate figure 1 through 4 are available on Github (https://github.com/bsieveri/sieveritz-2022-mt-prelimbic). Microscopy data will be made available upon request.

## Results

### Optogenetic inhibition of MT input to prelimbic cortex biased rats towards a high cost-high benefit option

We trained twenty-two 4- to 7-week-old male Sprague-Dawley rats on a benefit-benefit, cost-cost and cost-benefit decision-making task (Figure 2A; see methods section for details). At 9 weeks of age, twenty-two rats that had learned all three decision-making tasks received unilateral virus injections into MT. We injected 50-70 nl of either an adeno-associated virus expressing archaerhodopsin (AAV5-CAG-ArchT-GFP, n=10; ArchT rats) or a control virus (AAV5-CAG-GFP, n=12; controls). In addition, we implanted a short optical fiber attached to a 590 nm light emitting diode (LED fiber optic) through the contralateral hemisphere into ipsilateral prelimbic cortical layer 1 (Figure 2B). We confirmed that virus injections were primarily confined to MT (Figure 2C). Tips of LED fiber optics were located in prelimbic cortical layer 1 in close proximity to virus-expressing MT axon terminals (Figure 2C).

**Figure 2.**
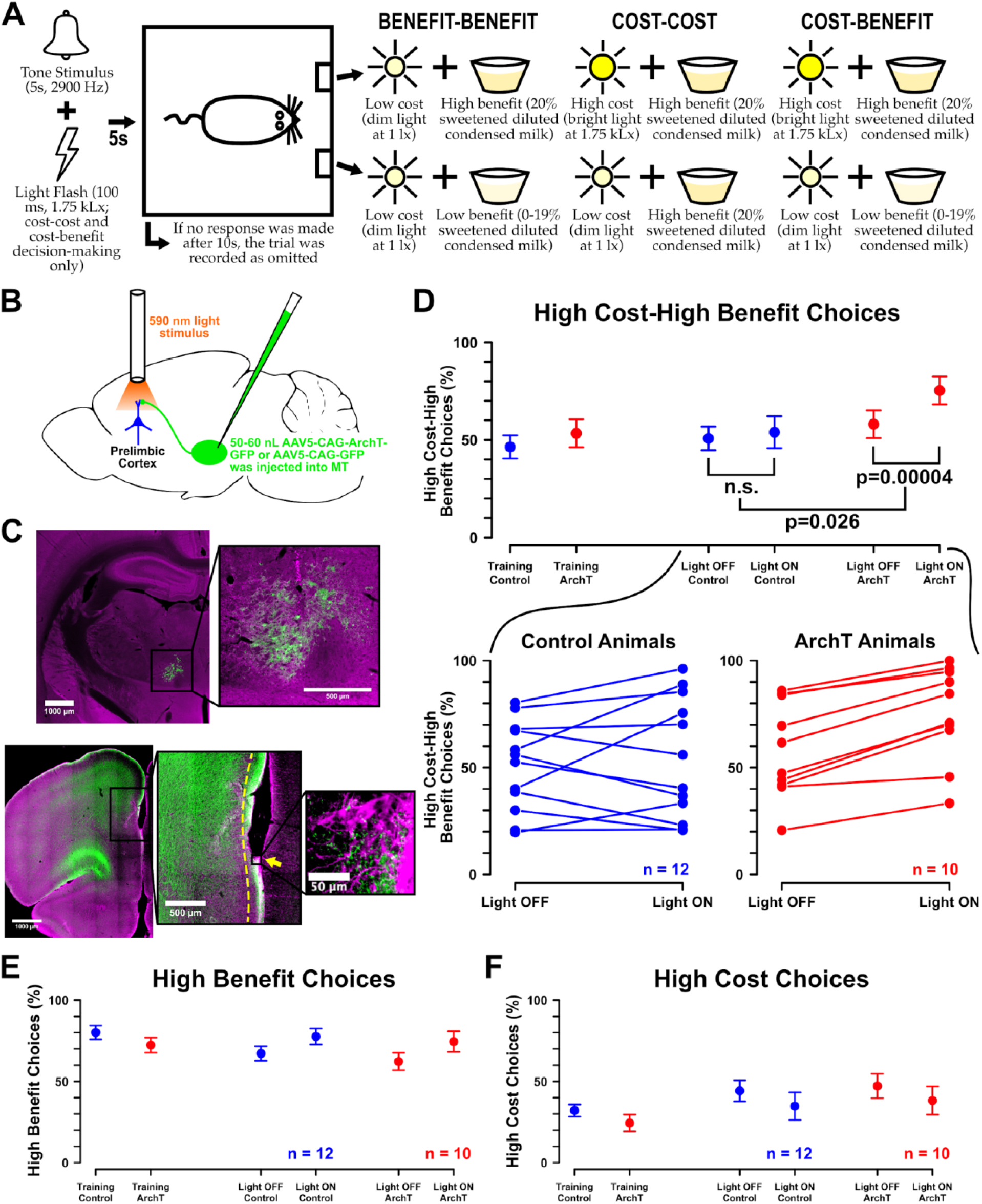
Optogenetic inhibition of MT axon terminals in prelimbic cortical layer 1 biases rats towards a high cost-high benefit option A-Illustration of the sequence of events on each trial. The dilution of sweetened condensed milk delivered as the low benefit was titrated, so that rats would choose both the high cost-high benefit and the low cost-low benefit option in about 50% of cost-benefit decision-making trials. B-50-70 nL of either AAV5-CAG-ArchT-GFP or the control virus AAV5-CAG-GFP were injected unilateral into MT and an LED fiber optic was implanted into ipsilateral prelimbic cortical layer 1. C-Upper illustration: Virus expression (green) was confined to MT, which was counterstained with GAD67 (magenta). Lower illustration: Virus-expressing MT axon terminals (green) and the LED fiber optic tip (yellow arrow) were located in prelimbic cortical layer 1. The yellow dashed line marks the approximate span of prelimbic cortical layer 1. Glial fibrillary acid protein, a neuronal marker for gliosis, is marked in magenta. D-Upper illustration: Mean percentage of high cost-high benefit choices +/- the standard error of the mean for controls (blue) and ArchT rats (red). For behavioral training, the mean across the first 20 trials on the last day of training is shown. For behavioral testing, data are separated between light ON and OFF trials. Lower illustration: Mean percentage of high cost-high benefit choices in light ON and OFF trials for each individual control (blue) and ArchT rat (red). E-Mean percentage of high benefit and F-high cost choices +/- the standard error of the mean for controls (blue) and ArchT rats (red).

**Figure 2 – figure supplemental 1.**
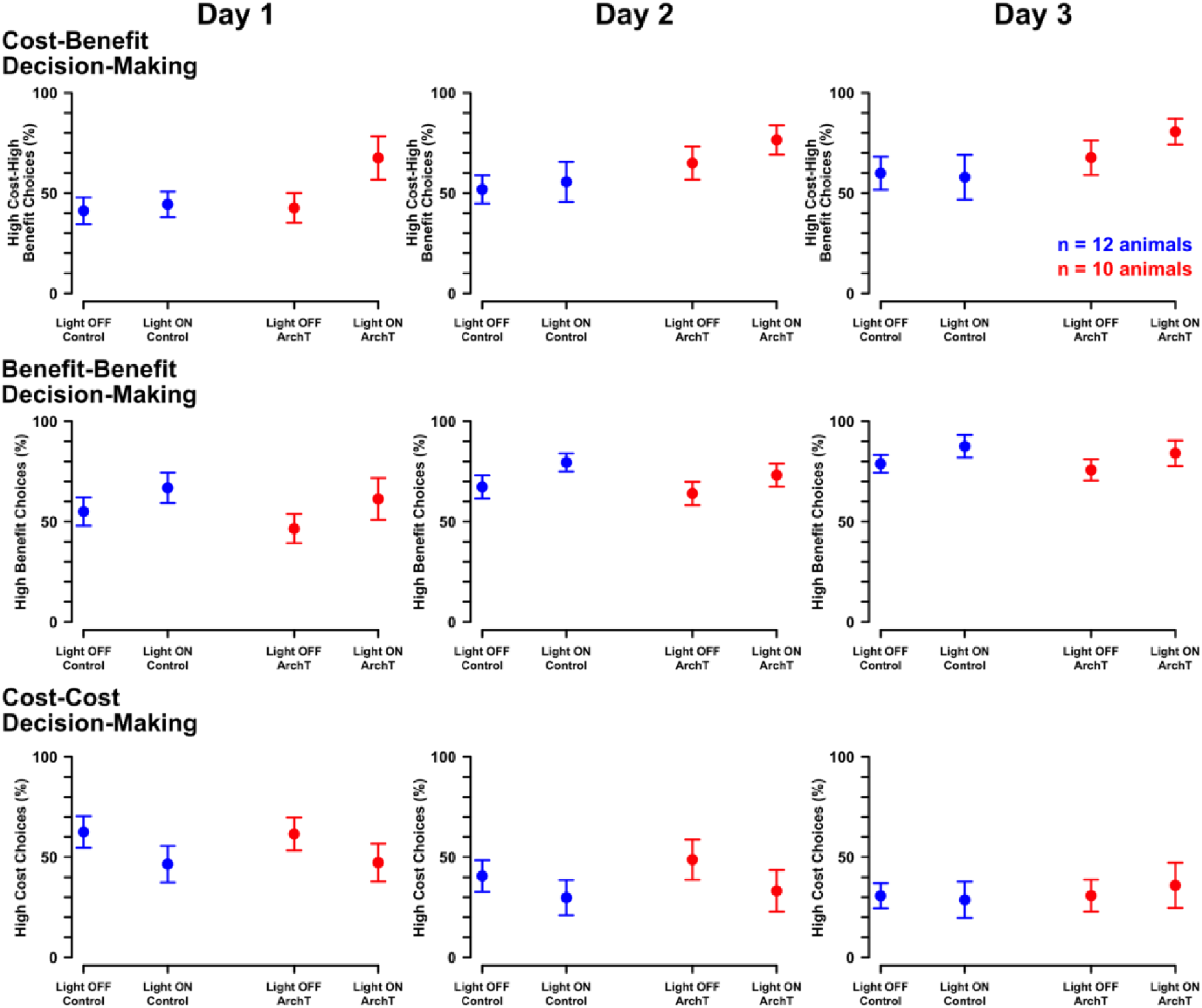
Data split by individual behavioral testing days showed similar trends to data averaged across all behavioral testing days. Mean percentage of high cost-high benefit, high benefit or high cost choices +/- standard error of the mean for controls (blue) and ArchT rats (red) in light ON and OFF trials on each individual day of behavioral testing.

**Figure 2 – figure supplemental 2.**
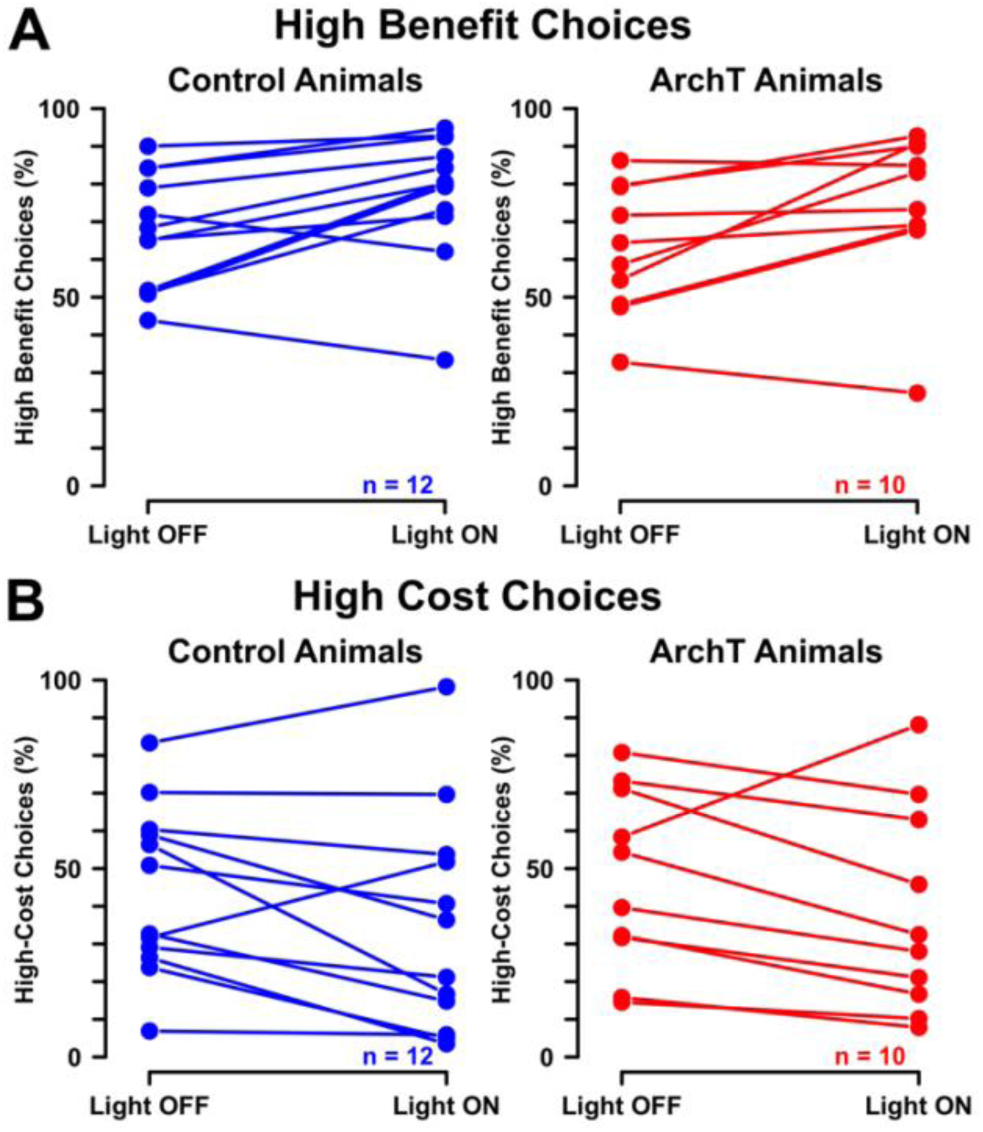
Data on the benefit-benefit and cost-cost decision-making task for each individual animal. A-Mean percentage of high benefit and B-high cost choices for each individual controls (blue) and ArchT rat (red) in light ON and OFF trials.

**Figure 2 – figure supplemental 3.**
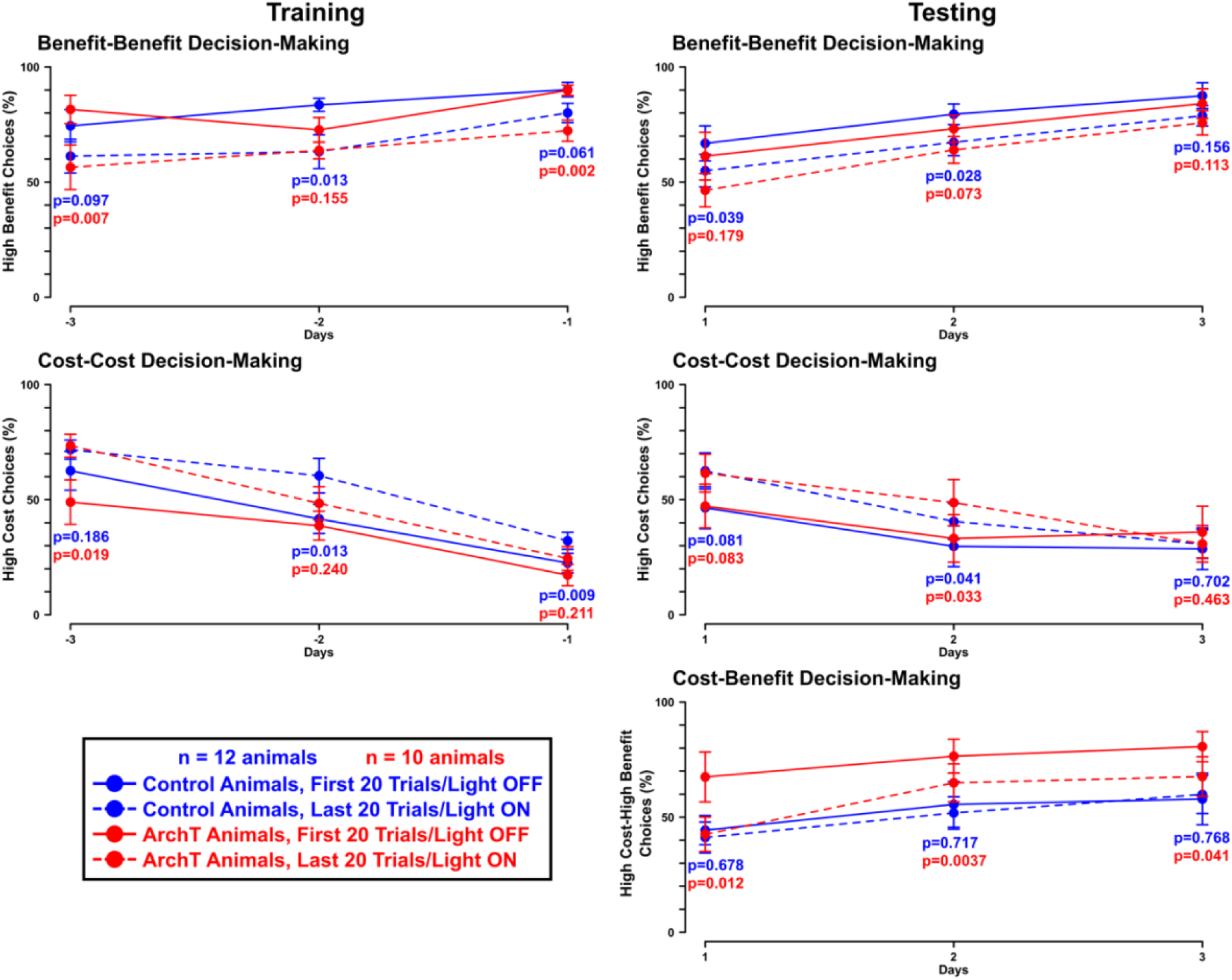
Rats show a within session retention effect on the benefit-benefit and cost-cost decision-making task. We compared the choice behavior on the first and last 20 trials on the last three days of behavioral training and on each day of behavioral testing. Plots show the mean percentage of high benefit, high cost or high cost-high benefit choices +/- the standard error of the mean for controls (blue) and ArchT rats (red) on the first or last 20 trials on each day on the last three days of behavioral training and on each day of behavioral testing. For behavioral testing the first 20 trials were light OFF trials, while the last 20 trials were light ON trials. We are not showing data on the percentage of high cost-high benefit choices for the last three days of behavioral training. Because the first three days of training on the high cost-high benefit decision-making task were used to titrate the dilution of sweetened condensed milk for each rat, data that includes them does not represent the actual choice behavior of rats after titration of the sweetened condensed milk dilution. Given that this is the case for most rats, we excluded the data from our analyses. Effects sizes are indicated in Appendix 1 – Table 1.

**Figure 2 - figure supplement 4.**
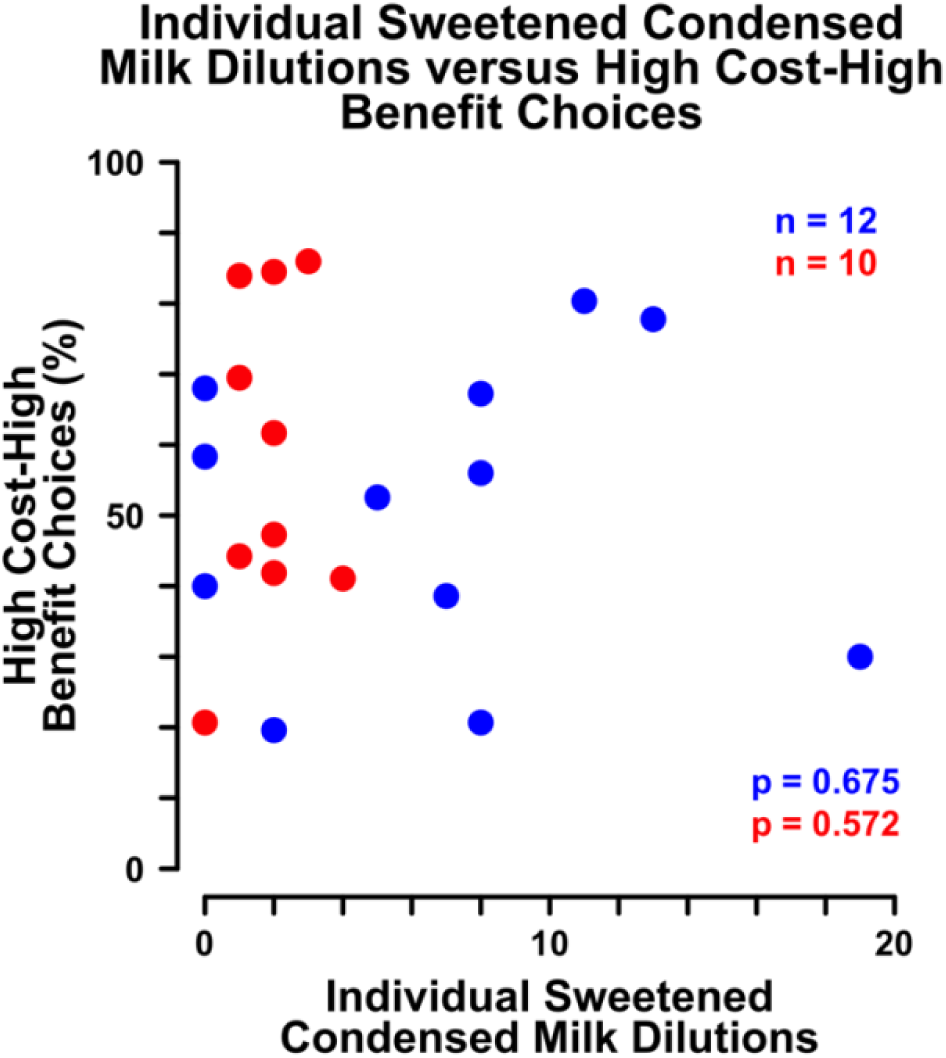
Correlation between individual dilution of sweetened condensed milk determined for each rat and high cost-high benefit choices for controls (blue) and ArchT rats (red).

**Figure 2 – figure supplemental 5.**
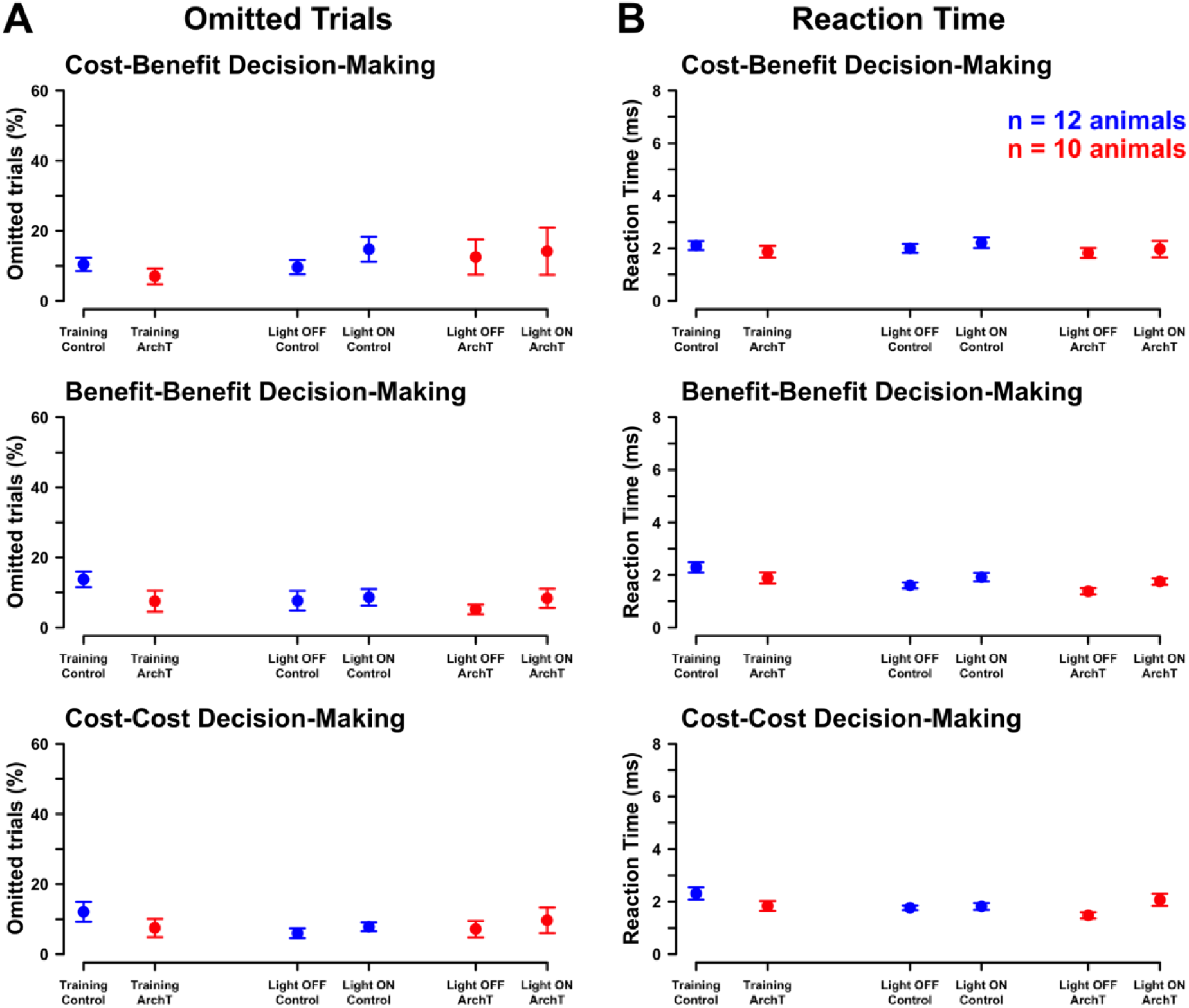
Percentage of omitted trials and reaction times on non-omitted trials upon optogenetic inhibition of MT axon terminals in prelimbic cortical layer 1. Plots show the mean percentage of omitted trials or reaction times on non-omitted trials +/- the standard error of the mean on the cost-benefit, benefit-benefit and cost-cost decision-making task for controls (blue) and ArchT rats (red) averaged over the first 20 trials on the last day of behavioral training, in light ON and OFF trials.

After a 12- to 16-day-long recovery period, we assessed rats’ choice behavior. For each of the three decision-making tasks, rats were presented with 40 trials per day on 3 consecutive days. We compared rats’ choice behavior without (light OFF) and with delivery of the 590 nm light (light ON). In ArchT rats, delivery of the 590 nm light induced optogenetic inhibition of virus-expressing MT axon terminals in prelimbic cortical layer 1. On each day, the first 20 trials were light OFF trials, while the last 20 trials were light ON trials. Optogenetic inhibition of MT axon terminals in prelimbic cortical layer 1 on the cost-benefit decision-making task biased rats towards the high cost-high benefit option (Figure 2D). In contrast, choice behavior on the benefit-benefit and cost-cost decision-making tasks was not affected (Figure 2E and 2F). To compare the percentage of high cost-high benefit choices on the cost-benefit decision-making task, we performed a mixed-design ANOVA with the injected virus as between animal factor and light ON/OFF as within animal factor. We observed a significant interaction effect between the injected virus and light ON/OFF (p=0.031, Cohen’s F=0.517, df=1). and a main effect for light ON/OFF (p=0.005, Cohen’s F=0.707, df=1). We performed post-hoc testing using multiple t-tests and applied a Bonferroni correction to account for multiple comparisons (adjusted significance level=0.0028). In light ON trials, we observed a significant increase in the percentage of high cost-high benefit choices for ArchT rats (p=0.00004, r=0.926, df=9), but not for controls (p=0.552, r=0.182, df=11). This indicates that optogenetic inhibition of MT axon terminals in prelimbic layer 1 biased rats’ choices towards the high cost-high benefit option (Figure 2D). To determine whether the increase in the percentage of high cost-high benefit choices from light OFF to light ON trials differed significantly between controls and ArchT rats, we used an uncorrected Student’s t-test. The increase was significantly larger for ArchT rats than for controls (p=0.026, r=0.536, df=20), further confirming that optogenetic inhibition of MT input to prelimbic cortex biased choices towards the high cost-high benefit option.

We observed a similar trend when data was split by day (Figure 2 - figure supplement 1). Given that only 20 light ON and OFF trials were administered on each individual day of behavioral testing, it is not surprising that none of the effects reached significance on any individual day. However, we did observe an increase in high-cost high-benefit choices in ArchT rats in light ON trials on the second day of behavioral testing that approached significance (p=0.0037, r=0.791, df=9; adjusted significance level=0.0028).

### No changes in choice behavior were observed on the benefit-benefit and cost-cost decision-making tasks

We observed a main effect of light ON/OFF in the benefit-benefit (p=0.001, Cohen’s F=0.903, df=1) and cost-cost decision-making tasks (p=0.014, Cohen’s F=0.605, df=1). However, post-hoc testing (Bonferroni correction applied to account for multiple comparisons; adjusted significance level=0.0028) did not confirm that the percentage of high benefit choices (ArchT: p=0.019, r=0.690, df=9; controls: p=0.016, r=652, df=11; Figure 2E and Figure 2 - figure supplement 2A) or high cost choices (ArchT: p=0.093, r=0.531, df=9; controls: p=0.076, r=0.508, df=11; Figure 2F and Figure 2 - figure supplement 2B) had changed significantly. This indicates that optogenetic inhibition of MT terminals in prelimbic cortex did not alter choice behavior in either of these two tasks.

Given that we used a block design and observed an increase in the percentage of high benefit choices and a decrease in the percentage of high cost choices from the first to the second half of each behavioral training session (Figure 2 – figure supplement 3), it is likely that the main effect of light ON/OFF was caused by a within session retention effect. However, the observed effect might instead have been due to unexpected electrophysiological side effects caused by exposure of the brain tissue to the 590 nm light, which was delivered to prelimbic cortical layer 1 in light ON trials. When we plotted the percentage of high benefit or high cost choices on the first and last 20 trials on the last 3 days of behavioral training (Figure 2 – figure supplement 3), we observed an increase of high benefit and a decrease of high cost choices between the first and last 20 trials on each of the last 3 days of behavioral training (Figure 2 – figure supplement 3). Given that we observed this effect during behavioral training, which happened prior to implantation of the 590 nm LED fiber optics, a within session retention effect is a more likely explanation than a side effect of the 590 nm light. In our experiments, the 590 nm light was delivered at a maximum intensity of 1.2mW and for a maximum of 20 secs in our experiments. In brain slices, delivery of a 532 nm light stimulus at 1 mW for up to 30 secs did not change the firing rate of cortical neurons (Stujenske et al., 2015), making it unlikely that light delivery alone increased or decreased neuronal activity. Moreover, given the numerical aperture and core radius of our optical fibers, a 561 nm light stimulus delivered at 1.2 mW is reduced to as little as 0.14 mW/mm^2^ at a 1 mm distance from the fiber tip (Deisseroth, 2021), making it even less likely that light delivery caused unexpected side effects in our experiments.

### Effects are not explained by pre-existing differences in choice behavior or caused by surgery

We confirmed that none of the behavioral effects, or the absence thereof, were explained by pre-existing differences in choice behavior. Choice behavior of controls and ArchT rats was comparable before and in light OFF trials after the surgery (p>0.0028; Figure 2D, 2E and 2F; for detailed p-values and effect sizes see Appendix 1 - Table 2). We further confirmed that behavioral effects or the absence thereof were not caused by surgery. No significant changes in choice behavior were observed pre- as compared to post-surgery for either controls or ArchT rats (p>0.0028; Figure 2D, 2E and 2F; for detailed p-values and effects sizes see Appendix 1 - Table 2). Lastly, we confirmed that reward sensitivity in light OFF trials as indicated by individually determined sweetened condensed milk dilutions did not correlate with the percentage of high cost-high benefit choices (Figure 2 - figure supplemental 4; ArchT: Kendall’s Tau=0.149, p=0.572, df=9; controls: Kendall’s Tau=0.095, p=0.675, df=11).

### Optogenetic inhibition of MT axon terminals in prelimbic cortex did not disrupt motor function

Given that we inhibited axon terminals emerging from MT, a brain region known to be heavily involved in motor control, we might have interfered with the rats’ motor function. To confirm that motor function was not disrupted, we compared the percentage of omitted trials as well as the average reaction time on non-omitted trials between light ON and OFF trials within ArchT rats and within controls (Figure 2 – figure supplement 5). Comparing the percentage of omitted trials, we observed a main effect for light ON/OFF on the cost-benefit decision-making task (p=0.035, Cohen’s F=0.506, df=1). Comparing the average reaction time, we observed an interaction effect (p=0.025, Cohen’s F=0.542, df=1) and a main effect for light ON/OFF on the cost-cost (p=0.013, Cohen’s F=0.607, df=1) as well as a main effect for light ON/OFF on the benefit-benefit decision-making task (p=0.001, Cohen’s F=0.855, df=1). However, post-hoc testing with an applied Bonferroni correction for multiple comparisons (adjusted significance level=0.0083) did not confirm any of these effects (p>0.0083, for detailed p-values and effect sizes see Appendix 1 - Table 2). Hence, we concluded that our rats’ motor function was not disrupted. Rats were able to move towards the lever and to move at comparable speeds in light ON and OFF trials.

### Optogenetic inhibition of MT input to prelimbic cortex decreases the activity of deep-layer pyramidal neurons

We explored a possible mechanism for prelimbic MT input to regulate cost-benefit decision-making. It has previously been shown that optogenetic inhibition of pyramidal neurons in prelimbic cortex biases rats towards a high cost-high benefit option (Friedman et al., 2015). We suggested that MT provides input to deep-layer neurons in prelimbic cortex, including pyramidal neurons, providing a mechanism for prelimbic MT input to regulate cost-benefit decision-making. In accordance with our first behavioral experiment, unilateral injections of 50-70 nl AAV5-CAG-ArchT-GFP were placed in MT in four 9- to 11-week-old male Sprague-Dawley rats. After a 12- to 16-day-long recovery period, rats were prepared for *in vivo* recording experiments under isoflurane anesthesia. An optical fiber, which was coupled to a 590 nm LED, was inserted through the contralateral hemisphere and placed in ipsilateral prelimbic cortical layer 1. To stimulate MT, a bipolar stimulating electrode was placed in ipsilateral MT. A glass electrode filled with 3.0M potassium methyl sulfate (KMeSO_4_) and goat anti-rat Alexa Fluor 594 (dilution between 1:50 to 1:200) was lowered into deep layers of ipsilateral prelimbic cortex to record extracellular neuronal activity (Figure 3A). Virus injection sites were comparable to the one presented in Figure 2B. Figure 3B and 3D illustrate the placement of optical fibers, bipolar stimulating electrodes and glass recording electrodes.

**Figure 3.**
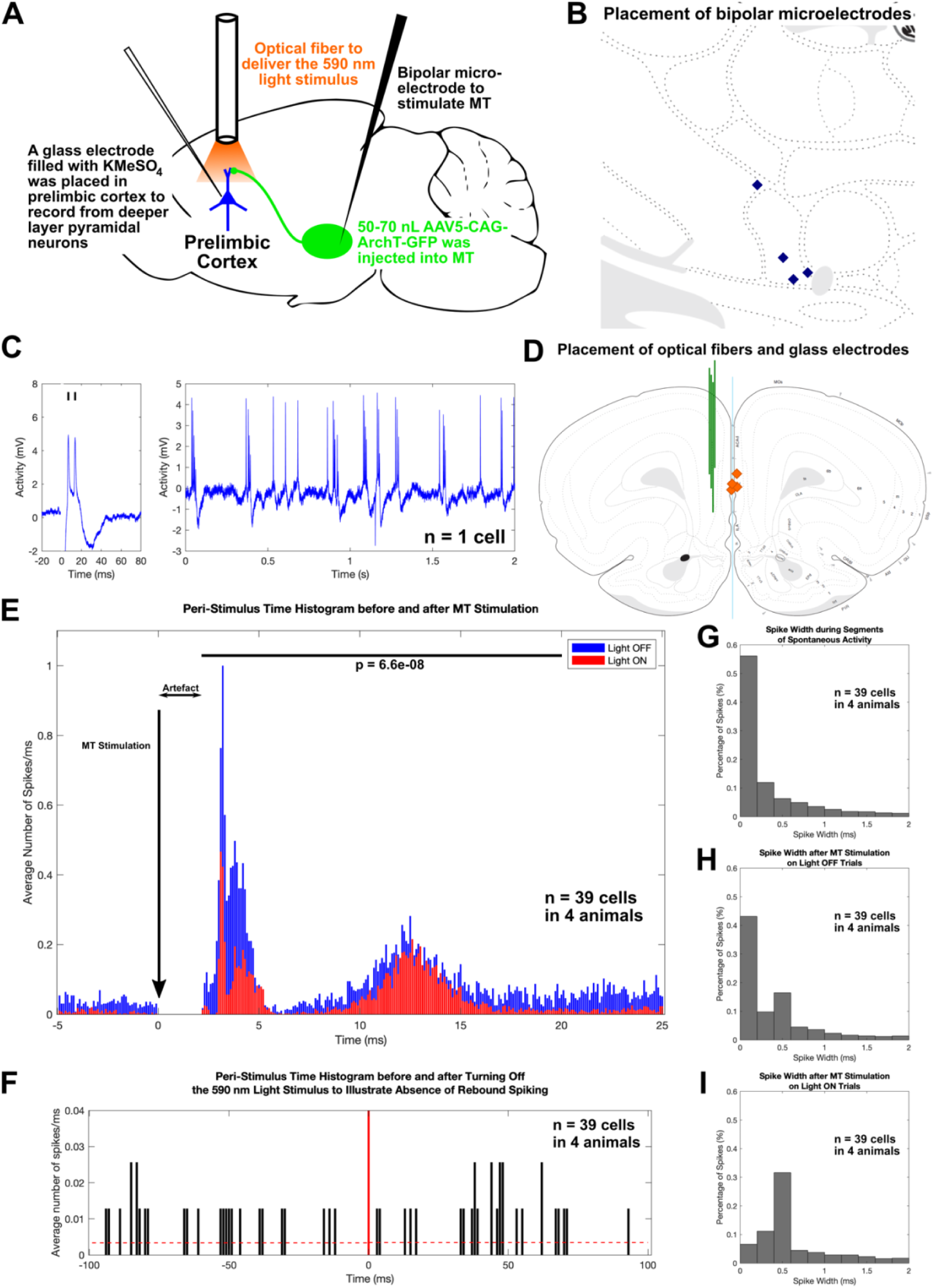
In anesthetized rats, MT stimulation induced a response in deep-layer prelimbic neurons and optogenetic Inhibition of MT input to prelimbic cortex reduced the response, A- MT was injected with 50-70 nL AAV5-CAG-ArchT-GFP. A glass electrode was used to record from neurons in deep layers of prelimbic cortex. MT was stimulated with a bipolar microelectrode. A 590 nm LED was placed in prelimbic cortical layer 1 to optogenetically inhibit MT axon terminals. B- Positioning of bipolar stimulating electrodes in MT. C- Neuronal response evoked by MT stimulation (left) and spontaneous activity recorded from one neuronal cluster (right). Extracellular spikes are marked by a small black line above the spike. D- Positioning of optical fiber tips (orange) and glass electrode tracks (green) in prelimbic cortex. E- Peri-stimulus time histograms show extracellular responses from 5 ms prior until 25 ms after MT stimulation in light OFF (blue) or ON trials (red). F- A peri-stimulus time histogram illustrates spontaneous neuronal activity in deep layers of prelimbic cortex. The vertical red line marks the time the 590 nm light was turned off. The red dotted lines indicate the mean number of spikes in the 100 ms before and after the light was turned off. G- Histogram of the width of spikes during segments of spontaneous activity, after MT stimulation on H-light OFF trials, or I-light ON trials.

While we recorded extracellular neuronal activity from neuronal clusters in deep layers of prelimbic cortex, we stimulated MT nuclei for 0.5 ms at −5 mA to induce a response in the recorded neuronal clusters (light OFF). Figure 3C shows an example of evoked spikes and of recorded spontaneous activity. When MT stimulation evoked a response, we simultaneously delivered 590 nm light to prelimbic cortical layer 1 to optogenetically inhibit MT axon terminals (light ON). Simultaneous optogenetic inhibition of MT input to prelimbic cortex did not completely abolish the response of neuronal clusters in deep layers of prelimbic cortex. However, the average number of extracellular spikes per millisecond observed between 2 and 20 ms after MT stimulation significantly decreased from an average of 0.11 spikes per millisecond without optogenetic inhibition to 0.06 spikes per millisecond under optogenetic inhibition (paired t-test, p=6.6e-8, df=38, Figure 3E). Our results indicate that MT input to neurons in deep layers of prelimbic cortex can evoke a response in these neurons. Optogenetic inhibition of MT input decreases the number of induced extracellular spikes. We propose that optogenetic inhibition of MT axon terminals in prelimbic layer 1 reduces the activity of corticostriatal pyramidal neurons in prelimbic cortex and, in turn, causes the observed behavioral bias towards a high cost-high benefit option.

Observed spikes are wider after MT stimulation than during segments of spontaneous activity (Figure 3G and 3H) and are widest when MT stimulation is paired with optogenetic inhibition of MT axon terminals in prelimbic cortical layer 1 (Figure 3I). Calcium-dependent spikes in cortex are wider (Stafstrom et al., 1985; Pockberger, 1991; De La Peña and Geijo-Barrientos, 2000), indicating that MT stimulation might induce calcium-dependent spikes. In addition, optogenetic inhibition of MT input to layer 1 seems to further increase the proportion of induced calcium-dependent spikes.

Although a previous study reported long-term effects upon sustained optogenetic inhibition of axon terminals such as an increase in spontaneous neurotransmitter release (Mahn et al., 2016), we did not observe any rebound spiking after optogenetic inhibition of MT axon terminals for as long as 20 secs. We never inhibited axon terminals for longer than 20 secs, indicating that the results outlined above were not caused by unexpected side effects from optogenetic inhibition (Figure 3F). Overall, our results are in line with previous studies that used the same virus (AAV5-CAG-ArchT-GFP) or the same viral construct as serotype 2 (AAV2-CAG-ArchT-GFP) to inhibit axon terminals, which showed robust behavioral effects (Stefanik and Kalivas, 2013; Stefanik et al., 2016) and partly confirmed that the virus is suitable for inhibiting axon terminals (Stefanik et al., 2013a, 2013b; Ozawa et al., 2017).

### MT input to prelimbic layer 1 inhibitory interneurons contributes to the regulation of neuronal activity in deep layers of prelimbic cortex

The ventromedial thalamus, one of the nuclei that make up MT, provides input to both prelimbic pyramidal neurons and prelimbic layer 1 inhibitory interneurons (Sieveritz and Arbuthnott, 2020). Hence, we were interested in whether prelimbic layer 1 inhibitory interneurons contribute to MT-mediated regulation of neuronal activity in deep layers of prelimbic cortex. We previously suggested that MT input regulates the activity of prelimbic layer 1 inhibitory interneurons. We further suggested that, in turn, layer 1 inhibitory interneurons regulate a network of cortical inhibitory interneurons, which ultimately regulates the activity of deep-layer pyramidal neurons (Sieveritz and Arbuthnott, 2020).

To test our prediction, we stimulated MT and simultaneously inhibited layer 1 inhibitory interneurons using optogenetics. In three 9- to 10-week-old Sprague-Dawley rats, we placed small unilateral injections of 10-20 nl AAV5-CAG-ArchT-GFP in prelimbic cortical layer 1. Given that prelimbic cortical layer 1 only contains inhibitory interneurons (Kubota, 2014) and no pyramidal neurons, this resulted in specific expression of AAV5-CAG-ArchT-GFP in prelimbic layer 1 inhibitory interneurons (Figure 4C). We observed no expression of the virus in pyramidal neurons or any other neurons in deeper layers of prelimbic cortex. Similar to the previous experiment, an optical fiber that was coupled to a 590 nm LED was inserted through the contralateral hemisphere and placed in ipsilateral prelimbic cortical layer 1. To stimulate MT, a bipolar stimulating electrode was placed in ipsilateral MT. A glass electrode filled with 3.0M potassium methyl sulfate (KMeSO_4_) and goat anti-rat Alexa Fluor 594 (dilution between 1:50 to 1:200) was lowered into deep layers of ipsilateral prelimbic cortex to record extracellular neuronal activity (Figure 4A). Figure 4B and 4D illustrate the placement of optical fibers, bipolar stimulating electrodes and glass recording electrodes.

**Figure 4.**
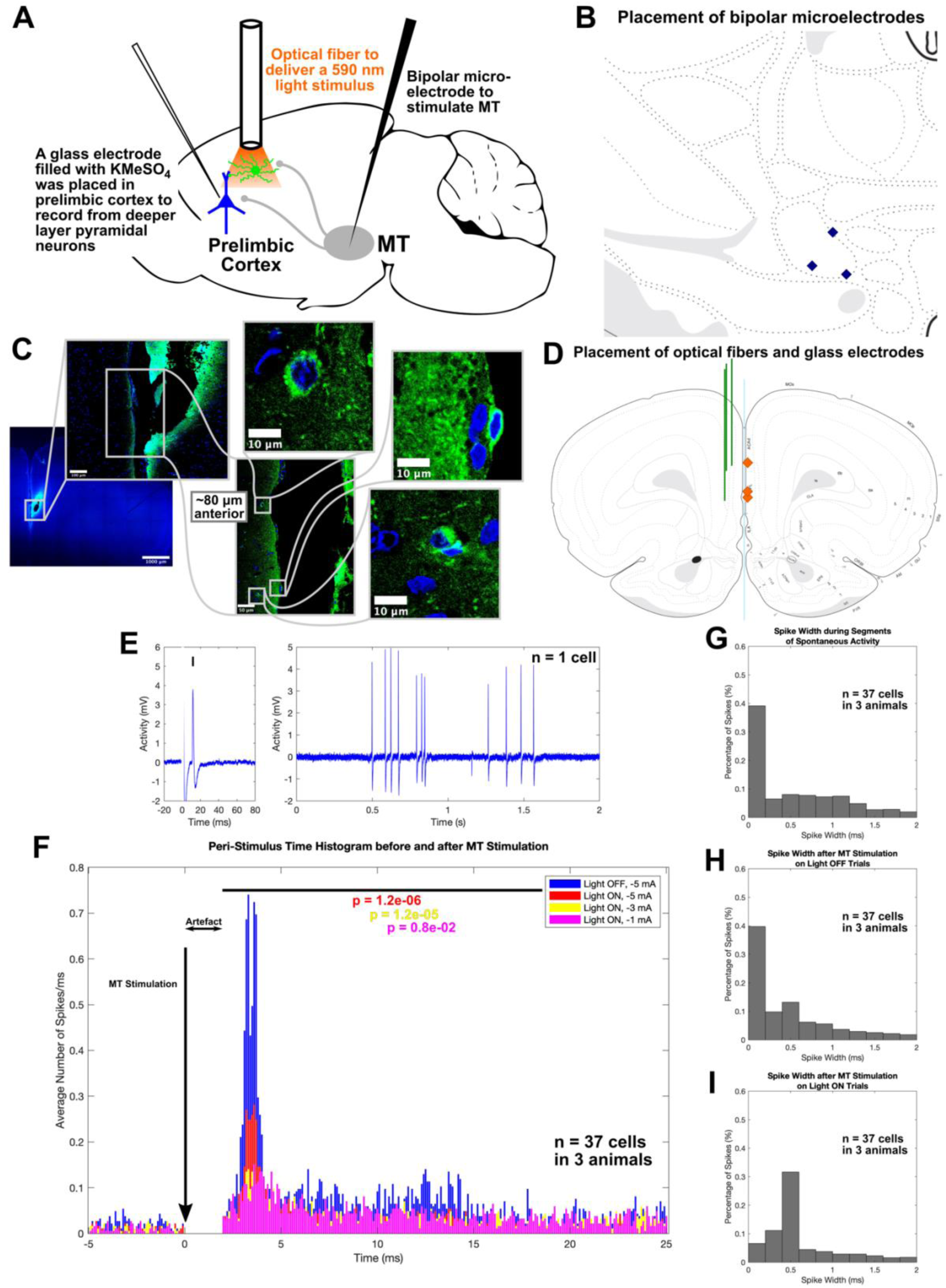
In anesthetized rats, optogenetic inhibition of layer 1 inhibitory interneurons in prelimbic cortex reduced extracellular responses to MT stimulation. A- Prelimbic cortical layer 1 was injected with 10-20 nL of AAV5-CAG-ArchT-GFP. A glass electrode was used to record from neurons in deep layers of prelimbic cortex. MT was stimulated with a bipolar microelectrode. A 590 nm LED was placed in prelimbic cortical layer 1 to optogenetically inhibit layer 1 inhibitory interneurons. B- Positioning of bipolar stimulating electrodes in MT. C- The optical fiber tip was located in and virus expression was limited to inhibitory interneurons (green) in prelimbic cortical layer 1. D- Positioning of optical fiber tips (orange) and glass electrode tracks (green). E- Neuronal response evoked by MT stimulation (left) and spontaneous activity recorded from the same neuronal cluster (right). Extracellular spikes are marked by a small black line above the spike. F- Peri-stimulus time histograms show extracellular responses extracellular responses in light OFF trials from 5 ms prior until 25 ms after MT stimulation at −5 mA (blue), and in light OFF trials after MT stimulation at −5 mA (red), −3 mA (yellow) and −1 mA (pink). G- Histogram of the width of spikes during segments of spontaneous activity, after MT stimulation on H- light OFF trials, or I- light ON trials.

We stimulated ipsilateral MT for 0.5 ms at −5 mA to induce responses in neuronal clusters in deep layers of prelimbic cortex (light OFF trials). Figure 4E shows an example of evoked spikes and of recorded spontaneous activity. When MT stimulation evoked a response, we simultaneously delivered 590 nm light to prelimbic cortical layer 1 to induce optogenetic inhibition in layer 1 inhibitory interneurons (light ON trials). Optogenetic inhibition of layer 1 inhibitory interneurons resulted in a decrease of evoked responses (Figure 4F). The average number of extracellular spikes per millisecond observed between 2 and 20 ms after MT stimulation decreased significantly; independent of the strength of MT stimulation (paired t-test; for MT stimulation of −5 mA: p=1.2e-6, df=36; for −3 mA: p=1.2e-5, df=36; for −1 mA: p=0.8e-2, df=36). Overall, our result indicates that prelimbic layer 1 inhibitory interneurons contribute to MT-mediated regulation of deep-layer neurons in prelimbic cortex.

Evoked spikes had comparable widths in segments of spontaneous activity and after MT stimulation (Figure 4G and 4H), but were wider when MT stimulation was paired with optogenetic inhibition of layer 1 inhibitory interneurons (Figure 4I). Given that calcium-dependent spikes in cortex are wider (Stafstrom et al., 1985; Pockberger, 1991; De La Peña and Geijo-Barrientos, 2000), inhibition of layer 1 inhibitory interneurons seems to increase the number of MT-induced calcium-dependent spikes. The result further underlines the importance of layer 1 inhibitory interneurons in processing of MT input to prelimbic cortex.

## Discussion

Optogenetic inhibition of MT axon terminals in prelimbic cortical layer 1 biased rats towards a high cost-high benefit option, indicating that MT input to prelimbic cortex contributes to cost-benefit decision-making. Rats were still able to accurately discriminate high and low benefit, and high and low cost options, indicating that MT input to prelimbic cortex is not necessary to process benefit or cost. However, it is crucial for trade-off choices that require rats to integrate multiple, conflicting reward values. When no integration of the two is required, inhibition of MT input to prelimbic cortex does not affect choice behavior. However, when integration is required, inhibition of MT input to prelimbic cortex biases rats towards a high cost-high benefit option. Our results demonstrate that MT input to prelimbic cortex is necessary to integrate multiple, conflicting reward values. We thus extend previous findings in mice that showed MT activity is correlated with action initiation (Gaidica et al., 2018) and reciprocal projections between MT and anterior lateral motor cortex play a role in sensory discrimination (Guo et al., 2017).

Integration of conflicting reward values might happen in MT before the outcome of the computation is transmitted to prelimbic cortex. Alternatively, MT input to prelimbic cortical layer 1 might modulate inputs from other brain regions to prelimbic cortex that drive integration of multiple, conflicting reward values. The latter explanation is supported by findings that MT input to prelimbic cortex is primarily modulatory (Collins et al., 2018). The former explanation highlights a long debate on whether thalamic nuclei only relay information between specific brain areas, or whether they perform computations on incoming streams of information and transmit the outcome of these computations to other brain areas. Research in the sensory domain demonstrated that sensory thalamic nuclei can integrate two incoming streams of signals (Saalmann and Kastner, 2011; Wolff et al., 2021). More recent research on the central thalamus, a region that borders with MT, has further shown that thalamic nuclei concerned with motor control might likewise integrate two streams of incoming signals (Matsuyama and Tanaka, 2021). In our case, MT might integrate benefit and cost before transmitting the outcome to prelimbic cortex.

We discovered a possible mechanism by which MT input to prelimbic cortex might regulate cost-benefit decision-making. Corticostriatal pyramidal neurons in deep layers of prelimbic cortex are known to be involved in cost-benefit decision-making (Friedman et al., 2015). We found that optogenetic inhibition of MT input to prelimbic cortex as well as inhibition of prelimbic layer 1 inhibitory interneurons reduced extracellular responses of neurons in deep layers of prelimbic cortex. We previously reported that about 80% of MT axon terminals that project onto prelimbic pyramidal neurons, project onto corticostriatal pyramidal neurons (Sieveritz and Arbuthnott, 2020). Given that MT stimulation evoked a response in the neuronal clusters that we recorded from, it is likely that we recorded from clusters of corticostriatal pyramidal neurons. Taking this assumption into consideration, we propose a two-fold mechanism to explain how MT input to prelimbic cortex regulates cost-benefit decision-making: 1) MT input directly regulates the activity of corticostriatal pyramidal neurons; and 2) MT input regulates the activity of layer 1 inhibitory interneurons, which induces feedforward inhibition in a network of cortical inhibitory interneurons. In turn, corticostriatal pyramidal neurons are disinhibited. When layer 1 inhibitory interneurons are inhibited, as was the case in our experiment, corticostriatal pyramidal neurons are no longer disinhibited and their activity is reduced. We further propose that the resulting decrease in corticostriatal pyramidal activity caused the observed bias towards a high cost-high benefit option (Figure 5). Such a mechanism is likely, given that direct optogenetic inhibition of corticostriatal pyramidal neurons similarly induced a bias towards a high cost-high benefit option (Friedman et al., 2015).

**Figure 5.**
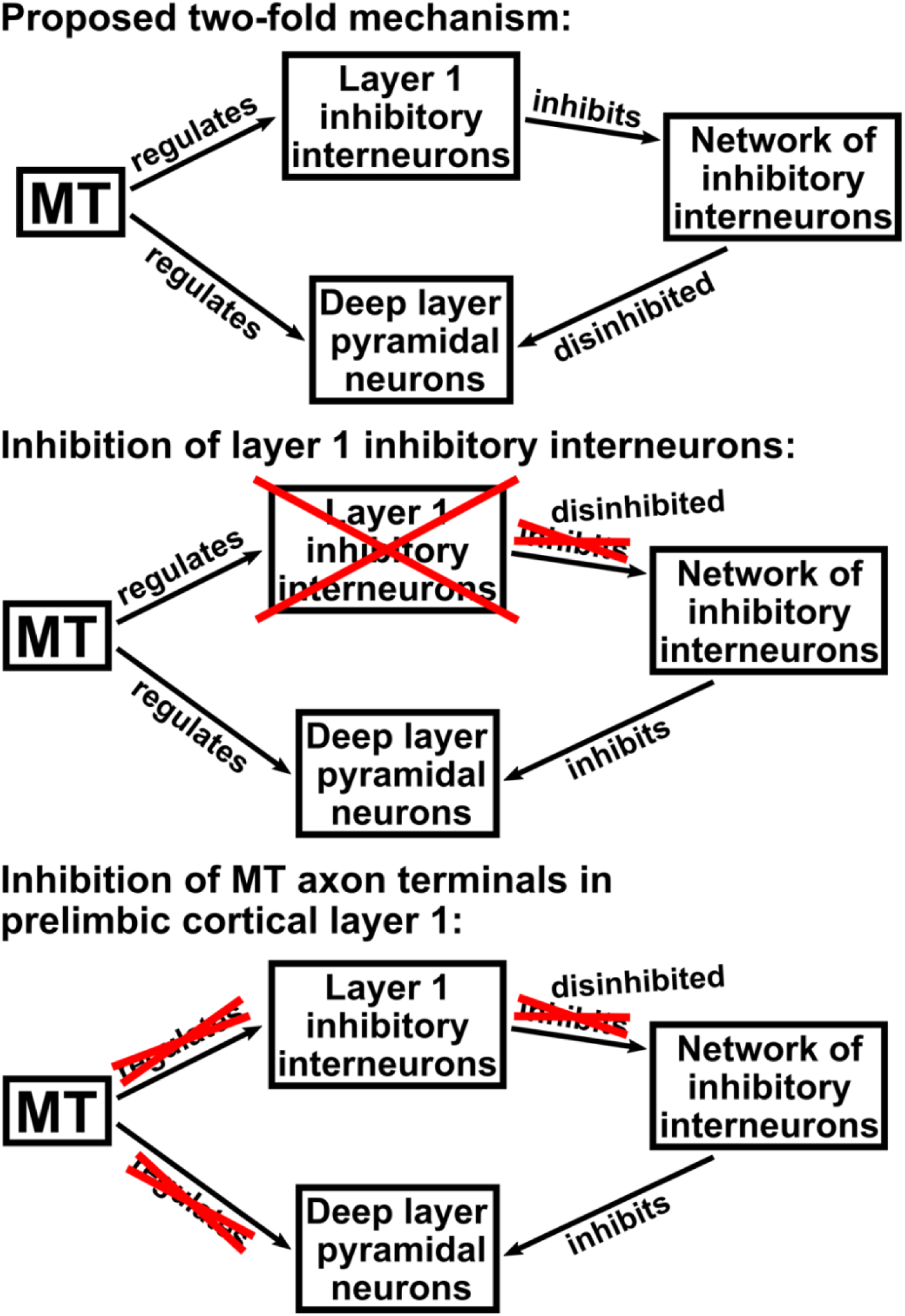
Proposed two-fold mechanism. MT input to prelimbic cortex regulates the activity of corticostriatal pyramidal neurons directly as well as indirectly via a network of cortical inhibitory interneurons. When layer 1 inhibitory interneurons are optogenetically inhibited, corticostriatal pyramidal neurons are no longer disinhibited and their activity is reduced. When MT axon terminals in prelimbic cortical layer 1 is optogenetically inhibited, the activity of corticostriatal pyramidal neurons is reduced due to missing MT input as well as due to them being inhibited by the network of cortical inhibitory interneurons. This causes the observed behavioral bias towards a high cost-high benefit option.

MT input might regulate the activity of corticostriatal pyramidal neurons through subthreshold excitation. When high negative currents that exceed physiological thresholds are used, as we did in this study, responses are evoked in deep-layer neuronal clusters in prelimbic cortex. In contrast, optogenetic stimulation of MT axon terminals in prelimbic cortex *in-vitro* induces subthreshold excitation in deep-layer pyramidal neurons, but does not evoke action potentials (Collins et al., 2018). Thus, rather than drive corticostriatal pyramidal neurons in prelimbic cortex, MT might regulate subthreshold excitation of these neurons by gating inputs from other subcortical sources to them. A similar mechanism was observed for the mediodorsal thalamic nucleus, which gates responses in prefrontal cortical neurons that were evoked by stimulation of the hippocampal output tract (Floresco and Grace, 2003). Alternatively, MT might regulate the activity of corticostriatal pyramidal neurons by long-term potentiation or depression, a mechanism that has been demonstrated for posterior medial thalamic input to barrel cortex (Williams and Holtmaat, 2019).

Aberrant decision-making is a symptom in anxiety (Aupperle and Paulus, 2010), depression (Amemori and Graybiel, 2012) and chronic stress (Schwabe and Wolf, 2009; Sousa and Almeida, 2012). Chronic stress biased rats towards high cost-high benefit options (Friedman et al., 2017), similar to what we observed when we inhibited MT axon terminals in prelimbic cortical layer 1. Given that optogenetic inhibition of MT input to prelimbic cortex emulates a behavioral phenotype observed under chronic stress, MT might play a crucial role in regulating stress. Patients with major depressive disorder exhibited aberrant task-related prefrontal cortical activity on an approach avoidance task that required integration of benefit and cost (Ironside et al., 2020). Hence, MT input to prefrontal cortex might also contribute to depression.

Our results show that MT contributes to choice behavior, especially in relation to trade-offs among conflicting reward values. These findings contrast with previous indications that MT is primarily associated with motor control. Further studies will be necessary to understand the full spectrum of MT contributions to cognition. Moreover, our findings suggest that, in addition to MT being involved in diseases related to motor control such as Parkinson’s disease (Brazhnik et al., 2016), MT might also be crucial in disorders associated with aberrant decision-making.

## Appendix 1

**Appendix 1 - Table 1.** Overview of p-values and effect sizes for t-tests comparing choice behavior on each individual behavioral testing day and on the last three days of behavioral training. The percentage of high benefit, high cost or high cost-high benefit choices within each group of rats was compared between light ON and OFF trials for each individual day of behavioral testing, or between the first and last 20 trials on the last three days of behavioral training. Training day −1 indicates the last day of behavioral training, training day −2 the second to last day and training day −3 the day prior to the second to last day. To account for multiple comparisons, t-tests were conducted using a Bonferroni corrected significance level of p=0.0028.

|  |  | Training |  |  |  |  |  | Testing |  |  |  |  |  |
| --- | --- | --- | --- | --- | --- | --- | --- | --- | --- | --- | --- | --- | --- |
|  |  | Day -3 |  | Day -2 |  | Day -1 |  | Day 1 |  | Day 2 |  | Day 3 |  |
|  |  | p-value | r (effect size) | p-value | r (effect size) | p-value | r (effect size) | p-value | r (effect size) | p-value | r (effect size) | p-value | r (effect size) |
| Benefit-Benefit Decision-Making | Change in High Benefit Choices (%), Controls | 0.097 | 0.48 | 0.013 | 0.668 | 0.061 | 0.533 | 0.039 | 0.575 | 0.028 | 0.606 | 0.156 | 0.418 |
|  | Change in High Benefit Choices (%), ArchT Rats | 0.007 | 0.757 | 0.156 | 0.459 | 0.002 | 0.821 | 0.179 | 0.437 | 0.073 | 0.561 | 0.113 | 0.505 |
| Cost-Cost Decision-Making | Change in High Cost Choices (%), Controls | 0.186 | 0.391 | 0.013 | 0.665 | 0.009 | 0.692 | 0.081 | 0.501 | 0.041 | 0.572 | 0.702 | 0.12 |
|  | Change in High Cost Choices (%), ArchT Rats | 0.019 | 0.687 | 0.24 | 0.387 | 0.211 | 0.409 | 0.083 | 0.545 | 0.033 | 0.642 | 0.463 | 0.248 |
| Cost-Benefit Decision-Making | Change in High Cost-High Benefit Choices (%), Controls | N/A | N/A | N/A | N/A | 0.306 | 0.455 | 0.678 | 0.127 | 0.717 | 0.111 | 0.768 | 0.091 |
|  | Change in High Cost-High Benefit Choices (%), ArchT Rats | N/A | N/A | N/A | N/A | 0.282 | 0.602 | 0.012 | 0.725 | 0.0037 | 0.791 | 0.041 | 0.623 |

**Appendix 1 - Table 2.** Overview of p-values and effect sizes for post hoc t-tests comparing the choice behavior, percentage of omitted trials and reaction times on non-omitted trials between ArchT rats and their controls as well as within each group pre- and post-surgery. Bonferroni-corrected significance levels for each metric are indicated in the bottom row.

|  |  | High cost-high benefit/high benefit/high cost choices (%) |  | Omitted Trials (%) |  | Overall Reaction Time |  |
| --- | --- | --- | --- | --- | --- | --- | --- |
|  |  | p-value | r (effect size) | p-value | r (effect size) | p-value | r (effect size) |
| Benefit-Benefit Decision-Making | ArchT Rats versus Controls, Pre-Surgery | 0.231 | 0.271 | 0.113 | 0.372 | 0.178 | 0.301 |
|  | ArchT Rats versus Controls, Post-Surgery | 0.493 | 0.161 | 0.444 | 0.195 | 0.189 | 0.294 |
|  | Pre- versus Post-surgery, Controls | 0.07 | 0.518 | 0.097 | 0.479 | 0.009 | 0.693 |
|  | Pre- versus Post-surgery, ArchT Rats | 0.011 | 0.505 | 0.437 | 0.262 | 0.017 | 0.698 |
| Cost-Cost Decision-Making | ArchT Rats versus Controls, Pre-Surgery | 0.243 | 0.282 | 0.25 | 0.256 | 0.132 | 0.333 |
|  | ArchT Rats versus Controls, Post-Surgery | 0.768 | 0.069 | 0.668 | 0.111 | 0.065 | 0.432 |
|  | Pre- versus Post-surgery, Controls | 0.043 | 0.569 | 0.019 | 0.637 | 0.047 | 0.56 |
|  | Pre- versus Post-surgery, ArchT Rats | 0.006 | 0.768 | 0.91 | 0.039 | 0.04 | 0.625 |
| Cost-Benefit Decision-Making | ArchT Rats versus Controls, Pre-Surgery | 0.463 | 0.172 | 0.261 | 0.26 | 0.403 | 0.2 |
|  | ArchT Rats versus Controls, Post-Surgery | 0.444 | 0.178 | 0.601 | 0.153 | 0.509 | 0.153 |
|  | Pre- versus Post-surgery, Controls | 0.589 | 0.165 | 0.731 | 0.106 | 0.063 | 0.53 |
|  | Pre- versus Post-surgery, ArchT Rats | 0.667 | 0.147 | 0.356 | 0.308 | 0.076 | 0.556 |
| Bonferroni corrected significance level: |  | p=0.0028 |  | p=0.0083 |  | p=0.0083 |  |

## Author Contributions

Conceptualization, B.S. and G.W.A.; Methodology, B.S., J.R.W. and G.W.A.; Software, B.S.; Formal Analysis, B.S.; Investigation, B.S. and S.H.D.; Writing – Original Draft, B.S., S.H.D., J.R.W. and G.W.A.; Visualization, B.S.; Supervision, J.R.W. and G.W.A.

## Notes

### Competing Interest Statement

The authors have declared no competing interest.

### Summary of Updates

Results on the absence of an effect on cost-benefit decision-making upon silencing motor thalamic input to striatum have been removed from the manuscript.

